# A mutation in *Hnrnph1* that decreases methamphetamine-induced reinforcement, reward, and dopamine release and increases synaptosomal hnRNP H and mitochondrial proteins

**DOI:** 10.1101/717728

**Authors:** Qiu T. Ruan, Neema Yazdani, Benjamin C. Blum, Jacob A. Beierle, Weiwei Lin, Michal A. Coelho, Elissa K. Fultz, Aidan F. Healy, John R. Shahin, Amarpreet K. Kandola, Kimberly P. Luttik, Karen Zheng, Nathaniel J. Smith, Justin Cheung, Farzad Mortazavi, Daniel J. Apicco, Durairaj Ragu Varman, Sammanda Ramamoorthy, Peter E. A. Ash, Douglas L. Rosene, Andrew Emili, Benjamin Wolozin, Karen K. Szumlinski, Camron D. Bryant

**Author notes:** Corresponding Author Camron D. Bryant, Ph.D., Department of Pharmacology and Experimental Therapeutics and Department of Psychiatry, 72 E. Concord St. L-616, Boston, MA 02118 USA, E. Equally contributing co-first authors (Qiu T. Ruan, Neema Yazdani). **Conflict of interest statement:** The authors declare no competing interests.

## Abstract

Individual variation in the addiction liability of amphetamines has a heritable genetic component. We previously identified *Hnrnph1* (heterogeneous nuclear ribonucleoprotein H1) as a quantitative trait gene underlying decreased methamphetamine-induced locomotor activity in mice. Here, mice (both male and female) with a heterozygous mutation in the first coding exon of *Hnrnph1* (H1+/-) showed reduced methamphetamine reinforcement and intake and dose-dependent changes in methamphetamine reward as measured via conditioned place preference. Furthermore, H1+/- mice showed a robust decrease in methamphetamine-induced dopamine release in the nucleus accumbens with no change in baseline extracellular dopamine, striatal whole tissue dopamine, dopamine transporter protein, or dopamine uptake. Immunohistochemical and immunoblot staining of midbrain dopaminergic neurons and their forebrain projections for tyrosine hydroxylase did not reveal any major changes in staining intensity, cell number, or in the number of forebrain puncta. Surprisingly, there was a two-fold increase in hnRNP H protein in the striatal synaptosome of H1+/- mice with no change in whole tissue levels. To gain insight into the molecular mechanisms linking increased synaptic hnRNP H with decreased methamphetamine-induced dopamine release and behaviors, synaptosomal proteomic analysis identified an increased baseline abundance of several mitochondrial complex I and V proteins that rapidly decreased at 30 min post-methamphetamine administration in H1+/- mice. In contrast, the much lower level of basal synaptosomal mitochondrial proteins in wild-type mice showed a rapid increase in response to methamphetamine. We conclude that H1+/- decreases methamphetamine–induced dopamine release, reward, and reinforcement and induces dynamic changes in basal and methamphetamine-induced synaptic mitochondrial function.

**SIGNIFICANCE STATEMENT:** Methamphetamine dependence is a significant public health concern with no FDA-approved treatment. We discovered a role for the RNA binding protein hnRNP H in methamphetamine reward and reinforcement. *Hnrnph1* mutation also blunted methamphetamine-induced dopamine release in the nucleus accumbens – a key neurochemical event contributing to methamphetamine addiction liability. Finally, *Hnrnph1* mutants showed a marked increase in basal level of synaptosomal hnRNP H and mitochondrial proteins that decreased in response to methamphetamine whereas wild-type mice showed a methamphetamine-induced increase in synaptosomal mitochondrial proteins. Thus, we identified a potential role for hnRNP H in basal and dynamic mitochondrial function that informs methamphetamine-induced cellular adaptations associated with reduced addiction liability.

## INTRODUCTION

Addiction to psychostimulants including methamphetamine (MA) is a major public health concern in the United States, with an estimated 535,000 individuals currently meeting the criteria for MA dependence (Lipari et al., 2016). Despite the prevalence of MA addiction, there is currently no FDA-approved treatment, in part because the neurobiological mechanisms underlying MA addiction are still not clear. Variation in sensitivity to the locomotor stimulant response to psychostimulants is a heritable trait and can sometimes predict differences in drug self-administration in rodents (Hooks et al., 1991; Yamamoto et al., 2013) because shared neurocircuits and neurochemical mechanisms underlie these behaviors. We recently used quantitative trait locus (QTL) mapping, positional cloning and gene editing via Transcription Activator-like Effector Nucleases (TALENs) to identify *Hnrnph1* as a quantitative trait gene for MA sensitivity in mice (Yazdani et al., 2015). *Hnrnph1* (heterogenous nuclear ribonucleoprotein H1) encodes an RNA-binding protein (RBP) that is expressed throughout the brain, and is a part of a subfamily of hnRNPs that includes hnRNP H1, hnRNP H2 and hnRNP F that possess structurally unique quasi-RNA recognition motifs (Honoré et al., 1995). hnRNP H1 regulates all aspects of RNA metabolism, including pre-mRNA splicing through binding at specific intronic sites, mRNA stability and translational regulation via binding to the 5’UTR and 3’UTR, and poly-adenylation control (Chou et al., 1999; George K. Arhin, Monika Boots, Paramjeet S. Bagga, 2002; Katz et al., 2010; Witten and Ule, 2011; Wang et al., 2012; Song et al., 2016).

We previously demonstrated that *Hnrnph1* polymorphisms and heterozygous deletion in the first coding exon of *Hnrnph1* affect behavioral sensitivity to acute MA-induced locomotor stimulation; however, the effects on MA reward and reinforcement are unknown. Additionally, the neurobiological mechanism(s) underlying the mutational effects of *Hnrnph1* on MA-induced behaviors remain to be established. *Hnrnph1* mRNA is ubiquitously expressed throughout the adult mouse brain (Lein et al., 2007). While the protein expression of hnRNP H1 appears to be nuclear-restricted, studies assessing hnRNP H1 protein in the brain are limited (Kamma et al., 1995; Honoré et al., 1999; Van Dusen et al., 2010). With regard to CNS function, hnRNP H family proteins are described as master regulators of neuron and oligodendrocyte differentiation via alternative splicing control (Wang et al., 2007; Grammatikakis et al., 2016). Whole-exome sequencing identified coding variants in human *HNRNPH1* and *HNRNPH2* (located on the X chromosome) associated with severe neurodevelopmental disorders (Bain et al., 2016; Pilch et al. 2018), implicating a crucial role of the hnRNP H protein family in neurodevelopment.

The purpose of the present study was three-fold. First, in order to expand beyond MA locomotor stimulant sensitivity, we examined the effect of the *Hnrnph1* mutation on oral MA reinforcement and intake via operant-conditioning and MA reward via conditioned place-preference (CPP). This mutation comprises a small, frameshift deletion in the first coding exon of *Hnrnph1* (H1+/-) that causes reduced MA-induced locomotor activity (Yazdani et al., 2015). To gain insight into the neurobiological mechanisms underlying behavioral differences in MA sensitivity, we examined drug-induced DA release via *in vivo* microdialysis, dopamine (DA) content of striatal tissue, and DA clearance from striatal tissue. Second, because we previously implicated *Hnrnph1* polymorphisms in dopaminergic neuron development, we assessed the effect of H1+/- on tyrosine hydroxylase (TH) levels in cell bodies and processes of the mesolimbic pathway via immunoblotting and immunohistochemistry. Finally, to gain insight into neural dysfunction in H1+/- mice at the protein level that could underlie behavioral and neurochemical deficits, we examined the synaptosomal proteome of the striatum between the H1+/- and wildtype (WT) mice at baseline and in response to MA.

## MATERIALS AND METHODS

### Mice

*Hnrnph1* mutant mice (H1+/-) were generated using TALENs targeting the first coding exon of *Hnrnph1* (exon 4, UCSC Genome Browser). Deletion of a small region in the first coding (exon 4) in *Hnrnph1* leads to a premature stop codon and transcription of a truncated mRNA message. Gene expression via qPCR with primers specific for exons 6 – 7 (not targeting the deleted exon 4) detected a 50% increase of *Hnrnph1* transcript in the H1+/- (heterozygous for deletion) mouse embryos relative to WT (Yazdani et al., 2015) and a 1.5-fold increase in the H1-/- (homozygous for deletion) (**Figure 7-1A to B**) with no change in gene expression of *Hnrnph2* (Yazdani et al., 2015; **Figure 7-1C**). In addition, gene expression via qPCR with primers specific for exon 4 detected a 50% reduction of *Hnrnph*1 transcript in H1+/- adult brain tissues relative to WT (Yazdani et al., 2015). In addition, no change in hnRNP H protein expression was detected in H1+/- and H1-/- embryonic stage brain tissue homogenate (**Figure 7-1D to G**). Because hnRNP H1 and H2 have 96% amino acid sequence homology, there is no commercially available antibody that can differentiate the two proteins. Quantitative analysis of protein peptide from a previously unpublished hnRNP H immunoprecipitation mass spectrometry study identified a decrease in the peptide encoded by the deleted region in exon 4 of *Hnrnph1* in H1+/- compared to the WT mice (**Figure 7-2 A to B**). No genotypic difference was observed in unique peptides associated with hnRNP H2 (**Figure 7-2 C to D**).

Experimental mice were generated by mating H1+/- males with C57BL/6J females purchased from The Jackson Laboratory in Bar Harbor, ME USA (for studies in Boston University) or in Sacramento, CA USA (for studies in UC Santa Barbara). Offspring were genotyped as previously described (Yazdani et al., 2015). Unless otherwise indicated, both female and male littermates (56-100 days old at the start of the experiment), were used in the studies. Mice were housed in same-sex groups of 3-5 in standard mouse cages and housed within ventilated racks under standard housing conditions. All procedures conducted in mice were approved by the Boston University, UC Santa Barbara and Virginia Commonwealth University Animal Care and Use Committees. All experiments were conducted in strict accordance with the Guide for the Care and Use of Laboratory Animals. Colony rooms were maintained on a 12:12 h light–dark cycle.

### Genotyping of H1+/- and H1-/- mice

The genotyping protocol for the H1+/- mice is published in Yazdani et al. (2015). For genotyping of H1-/- embryo tissue, an *Hnrnph1* forward primer (GATGATGCTGGGAGCAGAAG) and reverse primer (GGTCCAGAATGCACAGATTG) were designed to target upstream and downstream of the deleted region in exon 4 of *Hnrnph1*. Genomic DNA was used to amplify a 204-bp PCR product using DreamTaq Green PCR Mastermix (ThermoScientific) followed by overnight restriction enzyme digest with BstNI (New England Biolads).

### *Hnrnph1* and *Hnrnph2* RT-qPCR for mouse embryo tissue

Oligo-dT primers were used to synthesize cDNA from total RNA to examine mRNA expression and qPCR for evaluating gene expression were performed using Taqman SYBR Green (ThermoFisher Scientific Cat# 4309155). Each sample was run in triplicate and averaged. Differential gene expression was reported as the fold-change in H1+/- and H1-/- relative to WT littermates using the 2^-(ΔΔC_T_)^ method. The primer sequences used for evaluating expression of *Hnrnph1* (targeting exons 6 and 7 were ACGGCTTAGAGGACTCCCTTT and CGTACTCCTCCCCTGGAAGT. The primer sequences used for quantifying expression of *Hnrnph2* (targeting exons 1 and 2) were TAGCCGTTTGAGGGAAGAAG and CCCTGTTAGAGTTTCTTCCAGGTA. The house keeping gene used was *Hprt* (targeting exons 7 and 9) with following primer sequence: GCTGGTGAAAAGGACCTCT and CACAGGACTAGAACACCTGC.

### Oral MA self-administration

The procedures for MA operant-conditioning were similar to those recently described (Lominac et al., 2016). Testing was conducted in operant chambers equipped with 2 nose-poke holes, 2 cue lights, a tone generator, and a liquid receptacle for fluid reinforcement (Med Associates Inc.). Mice were not water-restricted at any point during oral MA self-administration procedures. The vehicle for MA was filtered tap water. Under a fixed-ratio 1 (FR1) schedule of reinforcement, mice were trained daily to self-administer MA during 1 h sessions where a single active nose poke response resulted in delivery of 20 μl of liquid MA into the receptacle, with a 20 s illumination of the cue light, and the sounding of the tone. During the 20 s period, further responding resulted in no programmed consequences. Inactive hole responses were recorded but had no consequences, serving to gauge the selectively of responding in the MA-reinforced hole. Mice were initially trained to nose-poke 80 mg/l MA, with the concentration of MA progressively increased over weeks (120, 160, 200, 300 and 400 mg/l MA; five days per dose). Upon the completion of each daily session, the volume of MA remaining in the receptacle was determined by pipetting and was subtracted from the volume delivered to calculate MA intake (Lominac et al., 2016).

### Conditioned place preference (CPP)

Mice were trained for 1 h each day in Plexiglas activity boxes within sound-attenuating chambers (40 cm length x 20 cm width x 45 cm tall; divided into two sides with different plastic floor textures for CPP). Mice were recorded from above using infrared cameras (Swan) and tracked with ANY-maze (RRID:SCR_014289). The CPP paradigm was described in Kirkpatrick and Bryant (2015).

On training Days 2-5, mice were injected with either saline (Days 2 & 4) or MA (Days 3 & 5; saline, 0.5 or 2 mg/kg, i.p.) and confined to either the saline- or MA-paired side for 1 h.

### Stereotaxic surgery

The procedures to implant indwelling microdialysis guide cannulae bilaterally over the NAc of mice were described previously (Lominac et al., 2014, 2016). Mice were anesthetized under 1.5–2% isoflurane with 4% oxygen as a carrier gas, mounted in a Kopf stereotaxic device with tooth and ear bars adapted for mice. The mouse skull was exposed, leveled, and holes were drilled based on coordinates from Bregma for the NAC (AP: +1.3 mm, ML: ±1 mm, DV: −2.3 mm), according to the mouse brain atlas of Paxinos and Franklin (2001). The guide cannulae were lowered bilaterally such that the tips of the cannulae were 2 mm above the NAc. The skull was then prepared for polymer resin application and the guide cannulae were secured to the skull with dental resin. Post-surgery, mice were injected subcutaneously (s.c.) with warm saline and 250 μl of 2.5 mg/ml Banamine (Henry Schein Animal Health) and allowed to recover on a heating pad. Post-operative care was provided for four days, during which mice were injected with 250 μl of 2.5 mg/mL Banamine s.c. daily for the first two days. Mice were allowed a minimum 1-week recovery prior to *in vivo* microdialysis assessments.

### *In vivo* microdialysis & HPLC analysis

Conventional microdialysis was conducted using a within-subjects design to examine saline and acute MA-induced DA release (0.5 or 2 mg/kg, i.p.), using procedures similar to those described previously (Lominac et al., 2014, 2016). Microdialysis probes were inserted unilaterally and perfused with artificial cerebrospinal fluid for 3 h (2 μl/min), allowing for neurotransmitter equilibration. For DA no net-flux analysis, DA was infused at 0, 2.5 nM, 5 nM, and 10 nM, and dialysate was collected in 20-min intervals for 1h/concentration. On a subsequent day, mice were probed on the contralateral side, and following the 3 h equilibration period and 1 h of baseline dialysate collection, mice were injected i.p. with either 0.5 or 2.0 mg/kg MA and dialysate was collected in 20-min intervals for 3 h post-injection. HPLC analysis of DA was conducted as described previously (Lominac et al., 2014). Cannulae placement was determined on Nissl-stained coronal sections and only mice exhibiting correct placement within the NAc were included in analyses.

### Behavioral test battery

#### Prepulse inhibition of acoustic startle

This test was employed to assess sensorimotor gating. The apparatus and procedures were identical to those previously described in Szumlinski et al. (2005). Six trial types were conducted: startle pulse (st110, 110 dB/40 ms), low prepulse stimulus alone (st74, 74 dB/20 ms), high prepulse stimulus alone (st90, 90 dB/20 ms), low or high prepulse stimulus given 100 ms. before the onset of the startle pulse (pp74 and pp90, respectively) and no acoustic stimulus (st0; only background noise). All trials were presented in a randomized order; st0, st110, pp74, and pp90 trials were given 10 times, whereas st74 and st90 were presented five times. Background noise in each chamber was 70 dB and the average inter-trial interval lasted 15 s.

#### Novel object test

To assess anxiety-like behavior, mice were placed into a rectangular box (9.25 x 17.75 x 8” high) containing one small, inedible object for 2 min. During that time, animals were allowed to explore and interact with the object. The number of contacts and time in contact (sec) with the novel object were video-recorded and tracked with ANY-maze tracking software (RRID:SCR_014289). The apparatus and procedures used were identical to those previously described in (Szumlinski et al., 2005).

#### Marble burying

The marble burying test was used to measure anxiety-like defensive burying (Njung’e and Handley, 1991). In our paradigm, 12 square glass pieces (2.5 cm^2^ × 1.25 cm high) were placed in the animals’ home cage, 6 at each end. The latency to start burying the marbles was determined by a blind observer using a stopwatch and the total number of marbles buried following a 20 min trial was recorded.

#### Light/dark shuttle box

The light/dark shuttle box test was employed to assess exploratory and anxiety-like behaviors. Mice were placed into a polycarbonate box (46 cm long × 24 cm high × 22 cm wide) wide containing distinct open (light) and closed (dark) environments for a 15 min trial. These two environments were separated by a central divider with an opening. Mice were first placed on the dark side and the latency to enter the light side, number of light-side entries, and total time spent in the light-side of the shuttle box were recorded using ANY-maze tracking software (RRID:SCR_014289). An increase in latency to enter the light, uncovered, side was interpreted as an index of anxiety-like behavior.

#### Porsolt swim test

To assess depressive-like behavior (Porsolt et al., 1977), mice were placed into a pool (30 cm in diameter; 45 cm high) filled with room-temperature water up to 35 cm and allowed to swim for a total of 6 min. Time immobile (s), immobile episodes, and immobile latency (sec) were video-recorded and tracked by ANY-maze tracking software (RRID:SCR_014289).

#### Accelerating rotarod

To assess motor coordination, mice were trained on the rotarod (IITC life science ROTO-ROD series) for a total of 10 trials over 3 days: 3 trials the first two days and 4 trials on the final day. The rotarod started at 4 RPM and accelerated to 40 RPM in 60 s. The time (s) it took a mouse to fall (physically falling or hanging off rotarod) was manually scored. Time on the rotarod was averaged across the total ten trials for each mouse.

### Bitter/Quinine Taste Sensitivity

H1+/- and WT mice were allowed continuous-access in the home cage to 4 sipper tubes containing 0 (filtered tap water), 0.1, 0.3 and 0.6 mg/ml quinine (Sigma-Aldrich). The quinine concentrations selected for study were based on Eastwood and Phillips (2014). The mice and bottles were weighed prior to initial presentation and the bottle was weighed every 24 h thereafter. The difference in bottle weight was used to determine the volume consumed from each solution over each 24 h period, the average intake from each solution, and the average total volume consumed.

### Quantification of baseline monoamine neurotransmitters from whole striatal tissue

Drug-naïve striatum were harvested from H1+/- and WT littermates and flash-frozen on dry ice. The dissected tissue was sent to Vanderbilt University Neurotransmitter Core for the quantification of monoamine neurotransmitters using HPLC wth electrochemical detection.

### DAT-mediated DA uptake

Saline or MA (2.0 mg/kg) was administered interperitoneally in a volume of 10 ml/kg. After 2 h post-administration (2 h was chosen based on the microdialysis results), mice were decapitated, and DAT-specific [^3^H]DA uptake from synaptosome preparations was conducted as described previously (Kivell et al., 2014). Mice were rapidly decapitated, and striatal regions were dissected from the brain and collected in 10 volumes (wt/vol) of prechilled 0.32 M sucrose buffer (0.32 M sucrose in 5 mM HEPES, pH 7.5). The striatal tissue was homogenized and centrifuged at 1000 x g for 15 min at 4°C. The supernatant was centrifuged at 12,000 x g for 20 min, and the pellet was suspended in 0.32 M sucrose buffer. Striatal synaptosomes (30 µg) were incubated in a total volume of 0.3 ml of Krebs-Ringer-HEPES (KRH) buffer consisting of 120 mM NaCl, 4.7 mM KCl, 2.2 mM CaCl_2_, 10 mM HEPES, 1.2 mM MgSO_4_, 1.2 mM KH_2_PO_4_, 5 mM Tris, 10 mM D-glucose, pH 7.4 containing 0.1 mM ascorbic acid, and 0.1 mM pargyline at 37°C for 10 min with or without DAT- specific blocker GBR12909 (50 nM). Following incubation, 5 nM [^3^H]DA (63.2 Ci/mmol-dihydroxyphenylethylamine [2,5,6,7,8-3H]; PerkinElmer) and further incubated for additional 5 min. Uptake of DA was terminated with the addition of 500 nM DAT blocker GBR12909. The samples were filtrated over 0.3% polyethylenimine coated GF-B filters on a Brandel Cell Harvester (Brandel Inc.), and washed rapidly with 5 ml cold PBS. Radioactivity bound to the filter was counted using a liquid scintillation counter. DAT mediated [^3^H]DA uptake was determined by subtracting total accumulation of [^3^H]DA (absence of GBR12909) and in the presence of GBR12909. Uptake assays were performed in triplicates.

### Immunohistochemistry

For immunohistochemistry (IHC), drug-naïve H1+/- and WT mice were anesthetized with pentobarbital, and transcardially perfused with phosphate buffered saline (PBS) followed by 4% paraformaldehyde in PBS at room temperature. Next, brains were dissected and processed for tyrosine hydroxylase (TH) 3-3’-diaminobenzidine (DAB) IHC and analysis as previously described (Burke et al., 1990; Hutson et al., 2011), or double immunofluorescent IHC for hnRNP H and TH colocalization. For DAB IHC, coronal slices were blocked with 4% normal goat serum and then incubated for 48 h at 4°C with anti-hnRNP H (1:50,000, Bethyl Cat# A300-511A, RRID:AB_203269) or tyrosine hydroxylase (TH) (1:500, Santa Cruz Cat# sc-14007, RRID:AB_671397), and processed for DAB staining and analyzed as previously described (Hutson et al., 2011). For co-staining studies with hnRNP H and TH, tissues were blocked with superblock (ThermoFisher Scientific Cat# 37515), and incubated with anti-hnRNP H (Santa Cruz Cat# sc-10042, RRID:AB_2295514) and TH (1:500, Santa Cruz Cat# sc-14007, RRID:AB_671397) for 48 hours at 4°C. Next, tissues were incubated with donkey anti-rabbit Alexa Fluor 488 (1:500, Molecular Probes Cat# A-21206, RRID:AB_141708) and donkey anti-goat Alexa Fluor 633 (1:500, Molecular Probes Cat# A-21082, RRID:AB_141493), washed, and then coated with ProLong Diamond Antifade Mountant (ThermoFisher Scientific Cat#P36961), mounted onto slides, and imaged on the Leica SPE Confocal microscope.

### TH puncta quantification in the striatum

Entire coronal slices of rostral, medial, and caudal striatum were imaged at 40x magnification using a Nikon Eclipse 600 microscope. A 225,000 µm^2^ grid was overlaid onto these images using Image J and every 3^rd^ field of view within the striatum was graded. Number of puncta within a field of view was graded in ImageJ by subtracting out background signal, creating binary images from these files, and then counting puncta meeting roundness and diameter criteria (roundness <0.6, diameter 1-45 µm). Averages puncta densities for ventral, dorsal, and total striatum were calculated. Grading of the puncta was performed in Image J. The image was duplicated into a 1,000,000-pixel area followed by brightness/contrast adjustment and background subtraction. The threshold was set to 106. A binary image was then generated for puncta measurement. Roundness was set to 0.6-1 and size was set to 5 – 200 pixels.

### Dissection of mouse brain regions: striatum and midbrain

Live, rapid decapitation was used to avoid the effects of anesthesia or CO_2_ asphyxiation on gene expression. Immediately followed live-decapitation using large, sharpened shears with an incision just posterior from the ears, the mouse brain was removed quickly with forceps and transfer to a cold surface. The striatum has a somewhat darker appearance than the surrounding cortex. To dissect the dorsal and ventral striatum, fine-tip forceps were used to separate the midline of the brain and then the cortex and hippocampus were removed to reveal the striatum. To dissect the midbrain, a razor blade was used to make a rostral cut where the cerebral aqueduct begins and another caudal cut just before the start of the cerebellum.

### Methamphetamine-induced locomotor activity followed by tissue harvesting

On Days 1 and 2, all mice received a saline injection (10 ml/kg, i.p.) and were recorded for locomotor activity in Plexiglas chambers (40cm length x 20 cm width x 45 cm height) for 1 h. On Day 3, mice receive either saline or MA (2 mg/kg, i.p.; Sigma Aldrich) and were recorded for locomotor activity for 30 min and the whole striatum was harvested as described above at 30 min post-injection. Whole striata (left and right sides) were flash frozen in ethanol/dry ice bath and stored at −80°C for long-term storage. Four cohort of animals were run in this behavioral paradigm for tissue collection: 1) hnRNP H immunoprecipitation followed by mass spec; 2) synaptosome mass spec and hnRNP H immunoblot; 3) validation studies for mitochondrial protein immunoblots; 4) measurement of MA concentration in MA-treated striatal tissues. The mice that were used for hnRNP H immunoprecipitation were all MA-treated on Day 3.

### Preparation of synaptosomes

Striatal tissue collection from saline- or MA-treated mice was performed as described above. The tissues were subsequently processed to obtain synaptosomes using a Percoll (Sigma Aldrich Cat# 1644) gradient fractionation method, which was adapted from Dunkley et al. (Dunkley et al., 2008). Whole striata (left and right hemisphere) were placed in 1 ml sucrose homogenization buffer (2 mM HEPES, pH7.4, 320 mM Sucrose, 50 mM EDTA, 20 mM DTT) supplemented with protease and phosphatase inhibitor (ThermoFisher Scientific Cat# 78440). The brain tissues were lightly homogenized using a handheld motorized pestle. The homogenate for each sample was then centrifuged for 2 min at 3000 rcf and the supernatant (S1) was collected. The pellet (P1) was then resuspended in 500 μl of sucrose homogenization buffer and re-spun for 3 min at 3000 rcf. The supernatant (S1’) was collected and combined with S1 and then centrifuged for 15 min at 9200 rcf. The supernatant S2 was then removed and the pellet (P2) was re-suspended in 500 μl of sucrose homogenization buffer and loaded onto a Percoll density gradient consisting of 23%, 10%, and 3% Percoll (1 ml each) in Polycarbonate centrifuge tubes (13 x 51 mm; Beckman Coulter Cat# 349622). The gradients were then centrifuged for 15 min at 18,700 rcf. The distinct band between the 10% and 23% Percoll was collected as the synaptosome. The synaptosome fraction was then washed to 5 mL with 1X HBM buffer, pH 7.4 (140 mM NaCl, 5 mM KCl, 5 mM NaHCO_3_, 1.2 mM NaH_2_PO_4_, 1 mM MgCl_2_-6H_2_O, 10 mM glucose, 10 mM HEPES) to dilute out the Percoll by centrifuging for 12 min at 18700 rcf. The pellet was then re-suspended in 100 μl of HBM buffer to yield the final synaptosme fraction and BCA assay was used to determine protein concentration. A total of 30 μg of synaptosome was loaded per sample for SDS-PAGE and Western blotting as described below.

### SDS-PAGE and Western Blot

Brain tissues were homogenized using hand-held homogenizer in RIPA buffer with Halt^TM^ Proteatase & Phosphotase inhibitor cocktail (ThermoFisher Scientific Cat# 78840). For each sample, 30 μg of protein was heated in a 70°C water bath for 10 min prior to loading into into a 4-15% Criterion TGX precast Midi protein gel (Bio-Rad) for SDS-PAGE followed by wet transfer to nitrocellulose membrane. The membrane was then blocked with 5% milk for 1 h and probed with primary antibodies. For evaluating TH expression in brain tissues, overnight incubuation of the membrane at 4°C with anti-TH (1:50,000, Santa Cruz Cat# sc-14007, RRID:AB_671397) was performed followed by 1 h incubation with donkey anti-rabbit HRP (1:10,00, Jackson ImmunoResearch Labs Cat# 711-035-152, RRID:AB_10015282). For evaluating hnRNP H protein expression in mouse embryo tissues and in striatal synaptosome, the following antibodies were used: hnRNP H (C-term: 1:50,000, Bethyl Cat# A300-511A, RRID:AB_203269; N-term: 1:50,000, Santa Cruz Cat# sc-10042, RRID:AB_2295514) followed by 1 h incubation with the appropriate secondary antibodies. For vadilation of mitochondrial protein expression following MA treatment, the following priamry antibodies were used: ATP5A1 (1:2000, Abcam Cat# ab14748, RRID:AB_301447); ATP5F1 (1:5000, Proteintech Cat# 15999-1-AP, RRID:AB_2258817); and NDUFS2 (1:5000; abcam ab192022). We used the following loading controls: anti-β-actin (1:20,000, Sigma-Aldrich Cat# A2228, RRID:AB_476697); anti-GAPDH (1:20,000; Millipore Cat# MAB374, RRID:AB_2107445), and PSD95 (1:10,000, Cell Signaling Technology Cat# 3450, RRID:AB_2292883).

Mouse brain tissue processing for SDS-PAGE and DAT immunblotting was modified from Staal et al., 2007. Briefly, tissue was triturated using a 20-22 gauge needle in RIPA buffer (10 mM Tris, pH 7.4, 150 mM, NaCl, 1 mM EDTA, 0.1% SDS, 1% Triton X-100) supplemented with Halt^TM^ Proteatase & Phosphotase inhibitor cocktail (ThermoFisher Scientific Cat# 78840). 30 μg of protein of each sample was allowed to rotate at room temperature prior to SDS-PAGE instead of heating the sample at high temperature. For evaluating DAT expression in whole striatal tissue and striatal synaptosome, we conducted overnight incubation anti-DAT (1:2000; Millipore Cat# MAB369, RRID:AB_2190413) followed by 1 h incubation with goat anti-rat (1:500, Jackson ImmunoResearch Labs Cat# 112-035-003, RRID:AB_2338128).

All processed membranes were imaged via enhanced chemiluminescence photo-detection. Densitometry analysis in Image J was used for quantification.

### hnRNP H immunoprecipitation

Following the three-day locomotor paradigm assessing acute locomotor sensitivity in H1+/- versus WT mice as described above, striata were dissected from the mice at 30 min post-injection of MA or saline. Striatum was dissected from WT or H1+/- mice and frozen on dry ice and stored at −80°C. Striatal tissues were then homogenized using a microcentrifuge pestle in ice-cold RIPA buffer (50 mM Tris-HCl, pH 6.8, 150 mM NaCl, 5 mM EDTA, 1% Triton X-100, 0.1% SDS, 0.5% sodium deoxycholate) supplemented with protease and phosphatase inhibitor cocktails (ThermoFisher Scientific Cat# 78840) and incubated overnight at 4°C with gentle agitation. Lysates were centrifuged at 10,000 rpm for 15 min at 4°C, and the supernatant fraction was saved for protein quantification via BCA assay. 1 mg of striatal lysates was pre-cleared for 1 h with 80 ul Protein G Sepharose coated beads (ThermoFisher Scientific Cat# 101243) and then centrifuged at 4°C for 5 min at 1000 rpm. The pre-cleared lysates (supernatant) were then incubated overnight with 10 μg of rabbit anti-hnRNP H (Bethyl Cat# A300-511A, RRID:AB_203269) or control normal rabbit IgG antibody (Millipore Cat# NI01-100UG, RRID:AB_490574). The next day, 80 μl of Protein G Sepharose coated beads were added to the antibody-lysate mixture, and incubated for an additional 2 h. The beads were then washed 4 times in 1 ml lysis buffer, resuspended and centrifuged for 1 min at 1000 rpm each time. The beads were eluted by adding 60 μl of non-reducing SDS buffer and heating at 95°C for 10 min. hnRNP H immunoprecipitates were eluted in non-reducing SDS buffer as described above. 50 μl of each sample was separated by SDS-PAGE at 100 V for 30 min (∼2 cm) on a Novex Bolt 4-12% Bis-Tris gel. The gel was then washed 3 times in deionized H_2_O and stained with Simply Blue Coomassie SafeStain (ThermoFisher Scientific Cat# LC6060). The gel was then cut at ∼160 kDa to exclude the prominent non-reduced IgG band. Individual gel lanes were then separately excised and stored in pre-washed microcentrifuge tubes at 4°C prior to shipping to the UMass Worchester Proteomics and Mass Spectrometry facility for analysis by OrbiTrap liquid chromatography tandem mass spectrometry.

### TMT Labeling, High pH reverse phase HPLC fraction, followed by LC-MS/MS

Following the three-day locomotor paradigm assessing acute locomotor sensitivity in H1+/- versus WT mice as described above, striata were dissected from the mice at 30 min post-injection. Synaptosomes were isolated following the procedure outlined above for proteomic characterization. Samples were resuspended in 8 M urea for 30 min, followed by the addition of 5 mM DTT for 1 h. Lodoacetamide was then add to the samples that were incubated in the dark for 30 min. The urea concentration was diluted below 1 M with the addition of 50 mM of ammonium bicarbonate. The samples were then digested with trypsin (50:1, protein to enzyme ratio) overnight at 37°C and terminated with the addition of formic acid to 1%. The samples were desalted with a C18 tip. Peptide was determined by Pierce Quantitative Colorimetric Assay (ThermoFisher Scientific Cat# 23275), then 100 μg of peptide was resuspended in 0.1 M triethylammonium bicarbonate (TEAB) and incubated with the TMT plex isobaric labeling for 1 h at room temperature. To quench the reaction, 5% hydroxylamine was added to each sample and incubated for 15 min. Each sample was combined at equal amount and cleaned with C18 tips. One mg of labeled peptides was fractioned using a Waters XBridge BEH C18 (3.5μm, 4.6 ×250mm) column on an Agilent 1100 HPLC system. 48 fractions were collected and combined to 12 fractions and then dried. A C18 Acclaim PepMap 100 pre-column (3μm, 100 Å, 75μm × 2cm) hyphenated to a PepMap RSLC C18 analytical column (2μm, 100 Å, 75μm × 50cm) was used to separate peptide mixture. LC-MS/MS analyses were completed using an EASY nLC 1200 system coupled to a Q Exactive HF-X mass spectrometer. Full MS spectra were collected at a resolution of 120,000 with an AGC of 3 × 10^6^ or maximum injection time of 50 ms and a scan range of 350 to 1650 m/z. MS2 scans were performed at 45,000 resolution and using 32% total normalized collision energy. Source ionization parameters were optimized with the spray voltage at 2.1 kV, dynamic seclusion was set to 45 s.

### Proteomics Data Analysis and Pathway Enrichment Results

The acquired data was searched by MaxQuant against the UniProt mouse proteome data base with standard settings. (fragment ion mass tolerance of 20ppm, maximum missed cleavage of 2, oxidation as variable modification, false discovery was 1%, only protein groups identified with at least 2 or more peptides). The intensity data were filtered and normalized using R (RRID:SCR_001905) and the LIMMA package (RRID:SCR_010943) was used for differential analysis with Genotype and Treatment as factors. A ranked list was generated from the analysis and the fgsea R package was used to perform pre-ranked analysis, with proteins filtered for absolute log_2_FC > 0.2 and p < 0.05. Enrichment results for the comparisons were visualized using Cytoscape (RRID:SCR_003032) and EnrichmentMap (RRID:SCR_016052), with nodes representing pathways and edges representing overlap genes. Pathways were clustered and annotated with themes using AutoAnnotate (Reimand et al., 2019).

### Experimental design and statistical analyses

We used an effect size (Cohen’s d = 0.9) based on genotypic differences in MA-induced locomotor activity at 30 min post-MA in our previously published data from H1+/- mice (Yazdani et al., 2015). We used G*Power 3 (Faul et al., 2007) to calculate a sample size of n = 16 per genotype required for 80% power (p < 0.05). Statistical details of the experiments can be found in the figure legends. Data are presented as means of replicates from each experiment ± SEM. For experiments in which two conditions were compared, an unpaired, two-tailed Student’s t-test was used to analyze the data. For experiment in which multiple factors were evaluated, ANOVA was used to calculate statistical significance as indicated in the figure legends. Statistical p value threshold for Student’s t test and ANOVA was set to 0.05. All statistical analyses were performed in R (RRID:SCR_001905).

## Data and Code Availability

All related data and materials are available upon request. The mass spectrometry proteomics data are available in MassIVE under proteome exchange accession number PXD014813.

## RESULTS

### Oral MA self-administration

We previously demonstrated a robust reduction in sensitivity to the locomotor stimulant response to MA in H1+/- mice (Yazdani et al., 2015). To more directly test MA-induced motivated behaviors, we investigated MA reinforcement and intake in H1+/- mice using an oral MA self-administration paradigm in C57BL/6J mice (Szumlinski et al., 2017). The study showed that mice allocated the majority of their nose-poking behavior toward the MA-reinforced hole, indicating drug reinforcement. H1+/- mice presented fewer active nose-pokes at the 160, 200, 300 and 400 mg/L MA doses compared to WT mice (Figure 1A), with no difference in non-specific, inactive nose-pokes across a range of MA concentrations (80-400 mg/L) (Figure 1B). Consistent with their lower level of MA reinforcement, H1+/- mice also consumed less MA (intake; mg/kg/day) at the 200, 300 and 400 mg/L concentrations (Figure 1C). These results indicate that the H1+/- mice are less motivated to work for the MA reinforcer. No sex differences or interactions were observed for any measure during self-administration testing. The genotypic differences in high-dose MA intake did not relate to bitter taste sensitivity as quinine intake in the home cage was equivalent between H1+/- and WT mice (**Figure 1-1**).

**Figure 1.**
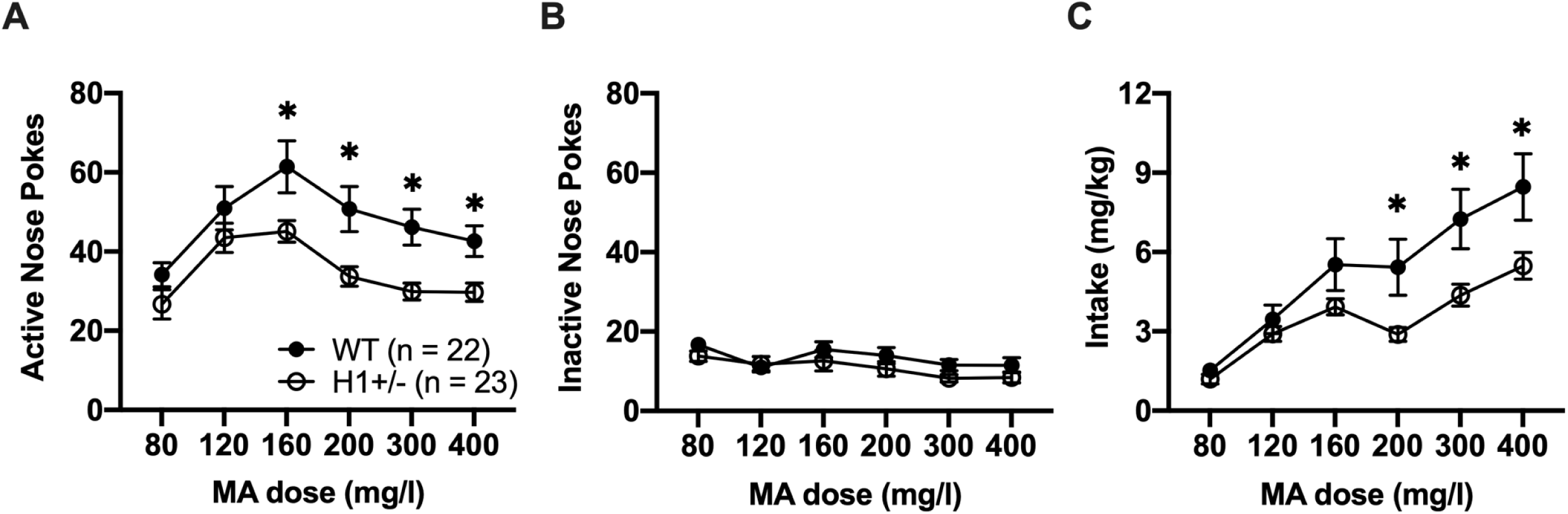
Oral MA self-administration in H1+/- mice. H1+/- mice were less sensitive than WT to the reinforcing effect of MA. H1+/- and WT mice were provided access to 80, 120, 160, 200, 300 and 400 mg/L of MA for a time period of five days per dose. **(A):** The average total active nose-pokes emitted during five, 1 h, sessions varied as a function of MA Dose [F(5,205) = 23.93, p < 0.0001] and Genotype [F(1,41) = 6.33; p = 0.02], with no Genotype x Dose interaction [F(5,205) = 1.582, p = 0.166] and no effect of Sex [F(1,41) < 1] with H1+/- showing less active nose pokes at MA doses 160, 200, 300, and 400 mg/l (main effect of Genotype at each dose: *p < 0.05). **(B):** The average total inactive nose-pokes emitted during the five, 1 h, sessions varied as a function of MA Dose [F(5,205) = 5.84, p < 0.0001], but not of Genotype [F(1,41) < 1], with no Genotype x Dose interaction [F(5,205) = 1.27, p = 0.28] or effect Sex [F(1,41) < 1]. **(C):** The average MA intake (mg/kg/day) exhibited by mice showed a significant Genotype x Dose interaction [F(5,205) =4.47, p < 0.0001], with H1+/- mice consuming less MA than WT mice at 200, 300, and 400mg/l doses of MA (main effect of Genotype at each dose: *p < 0.05). n = 23 (10 females, 13 males) for H1+/- and n = 22 (9 females, 13 males) for WT.

The blunted escalation of oral MA intake observed at high MA concentrations (≥ 200 mg/L) in H1+/- mice initially trained to respond for 80 mg/L MA prompted us to determine if the *Hnrnph1* mutation would blunt the acquisition of oral MA self-administration when 200 mg/L MA served as the reinforcer. However, we were unable to detect any genotypic difference in MA intake and active nose-pokes when mice were trained at this higher MA concentration (**Figure 1-2**). These results indicate that the *Hnrnph1* mutation interferes with the transition from low to high-dose MA self-administration, without altering the reinforcing effects of MA during acquisition.

### MA-conditioned reward

To further investigate why H1+/- mice showed less oral self-administration of MA, we assessed MA reward via conditioned place preference (CPP). When mice were tested in a MA-free state (Day 8), H1+/- mice showed lower CPP to 0.5 mg/kg MA but higher CPP to 2 mg/kg MA compared to the WT mice (Figure 2A). Similarly, during drug state-dependent CPP, H1+/- mice also exhibited lower and higher CPP at 0.5 mg/kg and 2 mg/kg MA doses, respectively (Figure 2B). No sex differences or interactions were observed for any measure during MA-CPP testing. The dose-dependent difference in CPP indicates that H1+/- showed a reduced sensitivity to the rewarding effect of MA where a higher dose of CPP was required to elicit CPP in H1+/- mice. In addition, although we did not detect significant genotypic differences in MA-induced locomotor activity during the confined one-half of the CPP box during training, we observed a trend toward a decrease in MA-induced locomotor activity in H1+/- mice on Day 9 during state-dependent CPP (following the two previous MA exposures) when they had twice as much open access space (**Figure 2-1**). This result was very similar to the previously published results in the same-sized open arena minus the CPP divider and a single MA exposure of 2 mg/kg (Yazdani et al., 2015). Furthermore, note that we replicated the reduced MA-induced locomotor activity phenotype in H1+/- mice with the same previously published protocol (Yazdani et al., 2015) prior to tissue harvesting for mass spectrometry analysis (**Figure 8-1**). There was no genotypic difference in the striatal level of either MA or its metabolite amphetamine at 30 min i.p. post injection of MA (**Figure 8-2**). Thus, differences in transport or metabolism of MA is an unlikely explanation for the dramatic genotypic differences in MA-induced behavior at 30 min post-MA.

**Figure 2.**
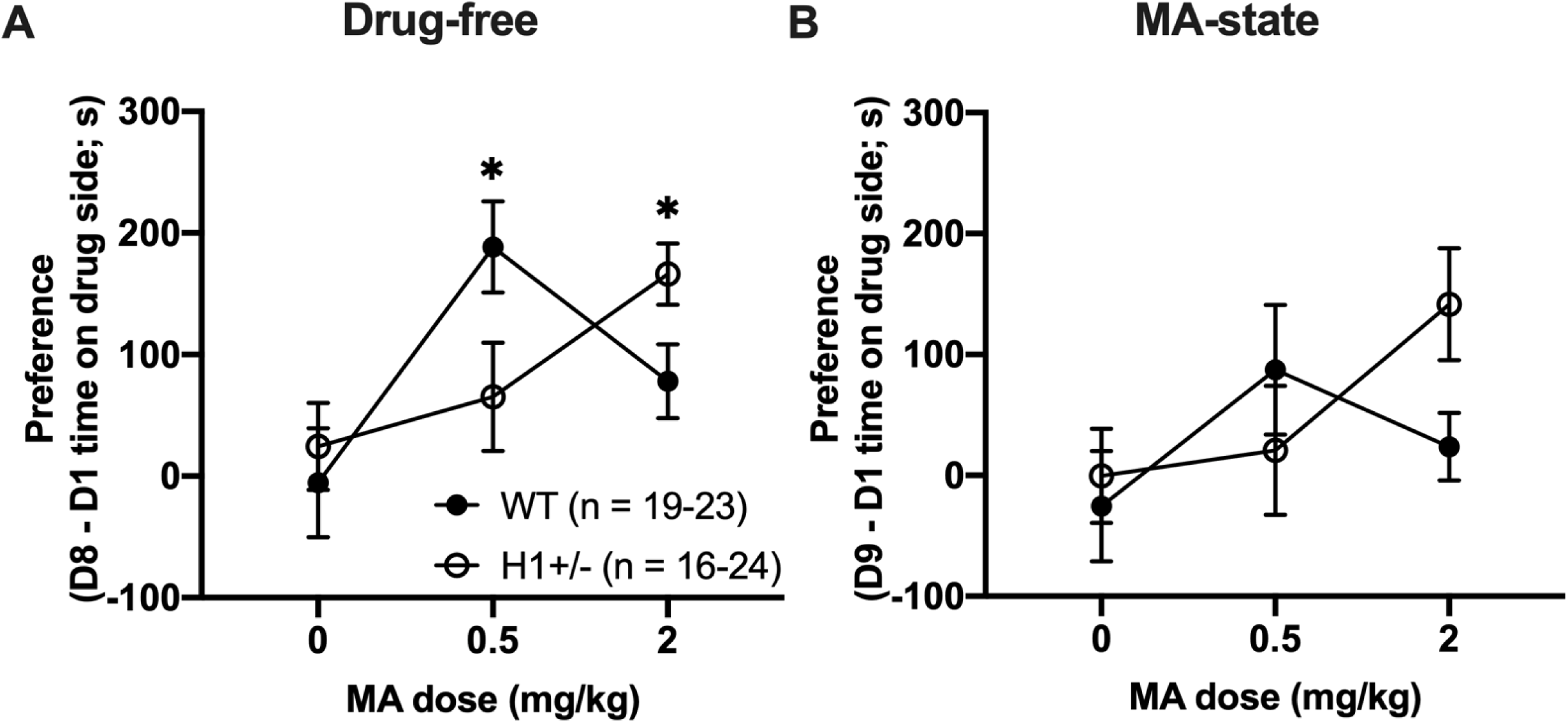
MA-induced CPP in H1+/- mice. H1+/- mice were less sensitive than WT to the rewarding effect of MA. **(A):** The genotypic difference in the time spent in the MA-paired side between Day 8 and 1 (preference in s) varied with MA doses [Genotype x Dose interaction: F(2,121) = 3.92, p = 0.023], with genotypic differences observed at 0.5 mg/kg [t(45) = −2.13, *p = 0.039] and 2 mg/kg MA doses [t(45) = 2.18, *p = 0.036]. No effect of Sex [F(1,115) = 0.77, p = 0.381] or Genotype x Sex interactions [F(1,115) = 0.04, p = 0.852] were observed. **(B):** In examining state-dependent CPP following a challenge dose of MA that was the same as during training, although a similar pattern of results was observed, there was no significant Genotype x Dose interaction [F(2,121) = 1.864, p = 0.16]. Furthermore, there was no effect of Sex [F(1,115) = 0.09, p = 0.761] or Genotype x Sex interactions [F(1,115) = 0.02, p = 0.882]. n = 24 (16 females, 8 males), 22 (14 females, 8 males) and 16 (9 female, 7 males) at 0, 0.5 and 2 mg/kg MA for H1+/-; n = 23 (9 female, 14 males), 23 (12 female, 11 males) and 19 (7 female, 12 males) at 0, 0.5 and 2 mg/kg MA for WT.

### MA-induced DA release in H1+/- mice

Because H1+/- mice showed reduced MA self-administration and reward, we next examined basal extracellular DA and MA-elicited DA release using *in vivo* microdialysis in the NAc, a brain region and neurochemical event that are necessary for MA reinforcement and reward (Prus et al., 2009; Keleta and Martinez, 2012; Bernheim et al., 2016). Linear regression analysis of no net-flux *in vivo* microdialysis in the NAc indicated no significant change in either DA release/reuptake (extraction fraction or slope, **Figure 3-1A**) or basal extracellular DA content or (**Figure 3-1B**). Following administration of 0.5 mg/kg MA (i.p.), WT mice exhibited an increase in extracellular DA whereas H1+/- mice showed a decrease in extracellular DA below baseline (Figure 3A). Administration of 2 mg/kg MA (i.p.) induced an increase in DA within the NAc of both genotypes, but H1+/- mice again showed markedly lower MA-induced extracellular DA levels, in particular from 100 to 180 min post-MA (Figure 3B). Consistent with the genotypic differences in extracellular DA levels, for both the 0.5 and 2 mg/kg MA doses, H1+/- mice also showed a lower level of 3,4-Dihydroxyphenylacetic acid (DOPAC, metabolite of DA) compared to WT mice (Figure 3C-D).

**Figure 3.**
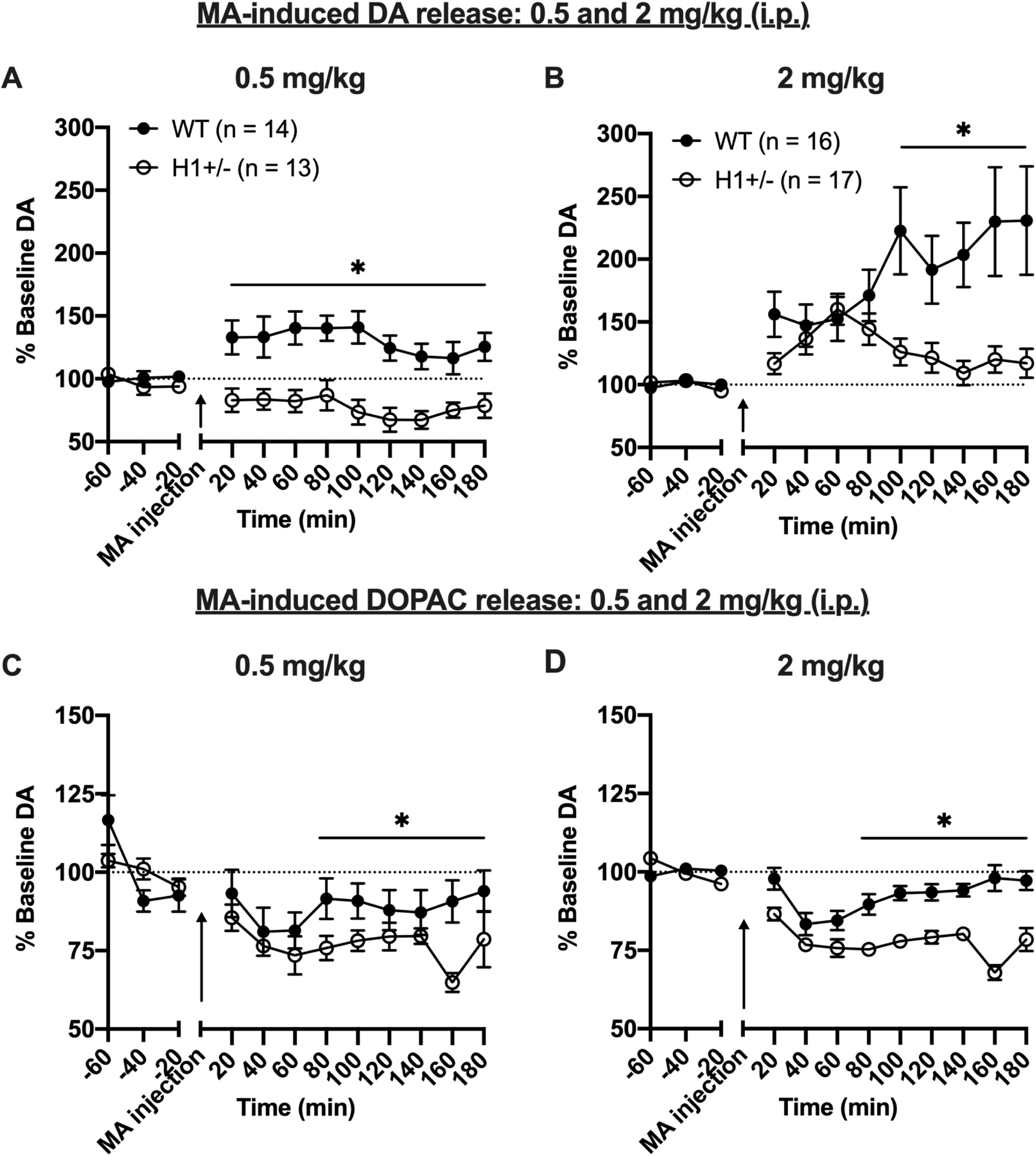
*In vivo* microdialysis of MA-induced DA release in H1+/- mice. H1+/- mice showed blunted MA-induced DA and DOPAC levels compared to WT. Mice were probed on the contralateral side, perfused with artificial cerebrospinal fluid (aCSF), and administered a MA challenge of either 0.5 mg/kg or 2 mg/kg (i.p., black arrow) after a 1 h baseline period. **(A-B):** The capacity of MA to elevate DA in the NAc was blunted in the H1+/- mice, in a manner that varied with the dose of MA administered [Genotype x Dose x Time: F(11,572)=2.31, p=0.01; no Sex effects or interactions, p > 0.25]. **(A):** The 0.5 mg/kg MA dose elicited an increase in extracellular DA in in WT but not in H1+/- mice [Genotype x Time: F(11,275) = 4.71, p < 0.0001], with H1+/- mice exhibiting significantly lower extracellular DA levels than WT mice at all time-points post-injection (main effect of Genotype: all *p’s < 0.04). **(B):** The 2 mg/kg MA dose elicited an increase in extracellular DA in both genotypes; however, the temporal patterning and the magnitude of this rise was distinct between H1+/- and WT mice [Genotype x Time: F(11,341) = 5.58, p < 0.0001]. H1+/- mice exhibited lower DA levels compared to WT mice during the first 20 min post-injection (main effect of Genotype: p = 0.05), as well as during the second half of testing [main effect of Genotype, *p’s < 0.025]. **(C-D)** MA dose-dependently reduced NAc extracellular DOPAC levels [Dose X Time: F(11,539) = 4.23, p < 0.0001]; however, irrespective of MA Dose or Sex, this reduction was, overall, greater in H1+/- versus WT mice [main effect of Genotype: F(1,49)=7.62, p = 0.008; Genotype X Time: F(11,539) = 5.61, p < 0.0001]. **(C)** 0.5 mg/kg MA induced a lowering of extracellular DOPAC, relative to baseline levels, with H1+/- mice exhibiting significantly lower DOPAC 80 min post-injection [Genotype X Time: F(11,308) = 2.78, p = 0.002; main effect of Genotype, *p = 0.03]. **(D):** 2 mg/kg MA induced a lowering of extracellular DOPAC in both genotypes; however, the effect was amplified in H1+/- mice [Genotype x Time: F(11,275) = 10.42, p < 0.0001]. Relative to WT animals, H1+/- mice exhibited lower DA levels during the first 20 min post-injection (main effect of Genotype: p = 0.05), as well as during the second half of testing [main effect of Genotype: *p’s < 0.025]. n=13 (7 females, 6 males) at 0.5 and n=17 (10 females, 7 males) at 2.0 mg/kg MA for H1+/-; n=14 (8 females, 6 males) at 0.5 mg/kg MA and 17 (10 females, 7 males) at 2 mg/kg MA for WT.

In examining glutamate levels, no net flux microdialysis study revealed no difference in basal extracellular level of glutamate (**Figure 3-2A**). Acute administration of 0.5 mg/kg or 2 mg/kg MA did not induce any effect on extracellular glutamate levels, nor was there any genotypic difference in glutamate levels (**Figures 3-2B and C**). To summarize, H1+/- mice showed a significantly blunted MA-induced DA release in the NAc in the absence of any significant difference in baseline DA levels and in the absence of any significant differences in glutamate neurotransmission. No sex differences or interactions were observed. These results point toward a MA-induced deficit in DA release as a plausible functional mechanism underlying the deficit in the MA-induced behavioral responses in H1+/- mice.

To further investigate the possible causes of the decreased capacity of MA to induce DA release in H1+/- mice, we quantified basal striatal tissue levels of biogenic amines in drug-naïve H1+/- and WT mice. No genotypic difference was detected in the amount of DA and DOPAC, or in the levels of other biogenic amines including HVA, 3-MT, 5-HT, etc. (Figure 4A). We also tested whether H1+/- impacts the function of DAT by examining DAT-mediated DA uptake in synaptosomes prepared from whole striatal tissue. However, no difference in basal DAT function was detected (Figure 4B). In addition, there was no change in the protein level of DAT in the total striatal brain lysate in H1+/- compared to WT mice (Figure 4C-D). There was also no effect of MA on DAT protein level in the synaptosome of either H1+/- or WT mice (**Figure 4-1).**

**Figure 4.**
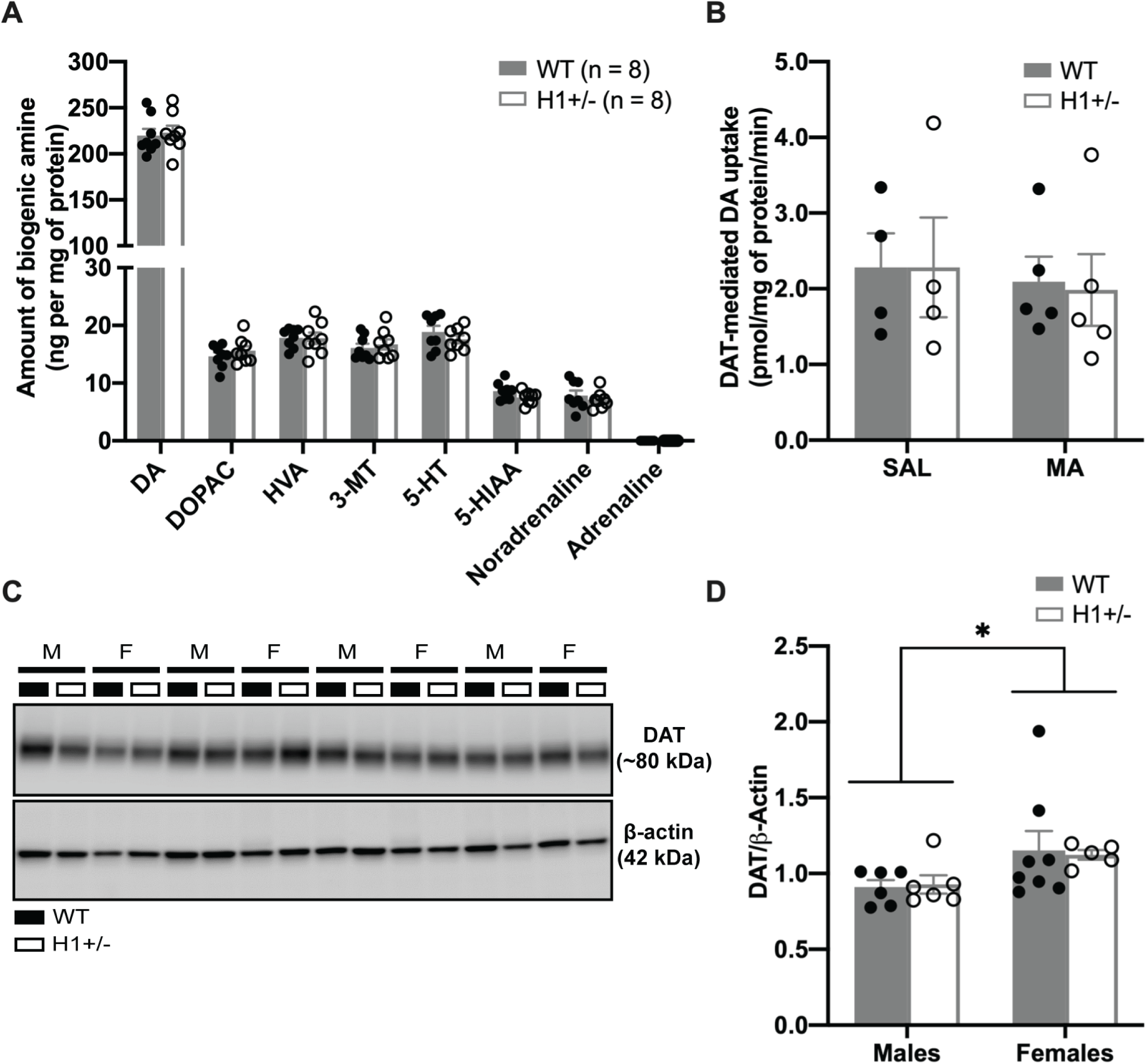
DA content and uptake and DAT levels in the striatum of H1+/- mice. No changes in DA or in DAT were detected at baseline in the striatum of H1+/- mice compared to WT mice. **(A)** Measurement of biogenic amine content in the striatum showed no difference in the level of dopamine between H1+/- and WT mice [WT, n = 8; H1+/-, n = 8; t(14) = −0.30, p = 0.8; unpaired Student’s T-test]. H1+/- also exhibited no change in the levels of other biogenic amines [p > 0.1; unpaired Student’s T-test]. **(B)** DAT-mediated DA uptake in the striatum of H1+/- mice. Striatum was harvested 2 h post-saline or MA injection for DA uptake assay, based on the time course of the microdialysis data in Figure 3B. There was no genotypic difference in the rate of DA uptake in response to saline (SAL) or MA treatment [Genotype x Treatment: F(1,14) = 0.02, p = 0.903]. **(C-D):** Immunoblot for DAT level in the striatum. Representative immunoblot for DAT shown in **(C)** with quantification shown in **(D)**. There was no change in DAT protein level in the striatum of H1+/- relative to WT mice [WT, n = 14; H1+/-, n = 11; t(23) = 0.32, p = 0.75; unpaired Student’s T-test]. There was a main effect of Sex where the female (regardless of Genotype) showed a higher level of DAT compared to that of male mice [female, n = 13; male, n = 12; t(23) = 2.53; *p = 0.02; unpaired Student’s T-test].

### Tyrosine hydroxylase expression in the mesolimbic and nigrostriatal brain regions of H1+/- mice

In our previous transcriptome analysis of congenic mice that captured *Hnrnph1* polymorphisms and decreased MA behavioral sensitivity, we identified a decrease in expression of *Nurr1* (*Nr4a2*), a transcription factor critical for dopaminergic neuron development (Yazdani et al., 2015). Thus, we hypothesized that dysfunctional mesolimbic and nigrostriatal DA pathways at the neuroanatomical level could underlie decreased MA-induced behaviors and DA release in H1+/- mice. We tested this hypothesis by examining changes in the number of neurons or the number of projections of neurons in the mesolimbic and nigrostriatal pathway, which mediate reward and motor activity, respectively (Adinoff 2004). Diagrams outlining the different brain regions assessed are presented in **Figure 5-1**. We first examined co-expression of hnRNP H (there are currently no commercially available antibodies for differentiating between hnRNP H1 and hnRNP H2) and tyrosine hydroxylase (TH; the rate-limiting enzyme for the synthesis of DA; Daubner et al., 2011). Results showed that hnRNP H and TH were co-expressed in the same midbrain TH neurons of the ventral tegmental area (VTA) and substantia nigra pars compacta (SNc) at a similar level in both genotypes (Figure 5A). hnRNP H immunostaining was primarily nuclear whereas TH immunostaining was cytoplasmic. We next examined expression of TH in the dopaminergic neurons of the mesolimbic and nigrostriatal circuit (Figures 5 and 6). TH DAB immunohistostaining in the VTA and SNc of the midbrain (Figure 5B) indicated no significant genotypic differences in TH optical density (OD) (Figure 5C). There was also no genotypic difference in the number (Figure 5D) or diameter (Figure 5E) of TH-positive neuron of the VTA and SNc. TH DAB immunostaining in the striatum indicated no change in TH OD in the dorsal striatum (Figures 6A-C) between H1+/- and WT mice. However, a small, but statistically significant increase in TH OD was observed in the ventral striatum (which includes the NAc) of the H1+/- mice (Figures 6D-F). To measure TH-positive puncta as an indirect estimate of the number of DA terminals in the striatum, we performed stereology under higher magnification (**Figure 6-1A**) and detected no difference in the number of TH-positive puncta between the H1+/- and WT mice in the dorsal striatum or in the ventral striatum (**Figure 6-1B**). Immunoblot analysis indicated no significant difference in TH protein level in either the midbrain (Figures 5F-G) or the whole striatum (Figures 6G-H). Taken together, analysis of the dopaminergic mesolimbic and nigrostriatal DA circuit did not provide strong evidence for changes in the expression of TH within cell bodies or terminals or changes in the number of TH-positive terminals that could explain behavioral differences in H1+/- mice.

**Figure 5.**
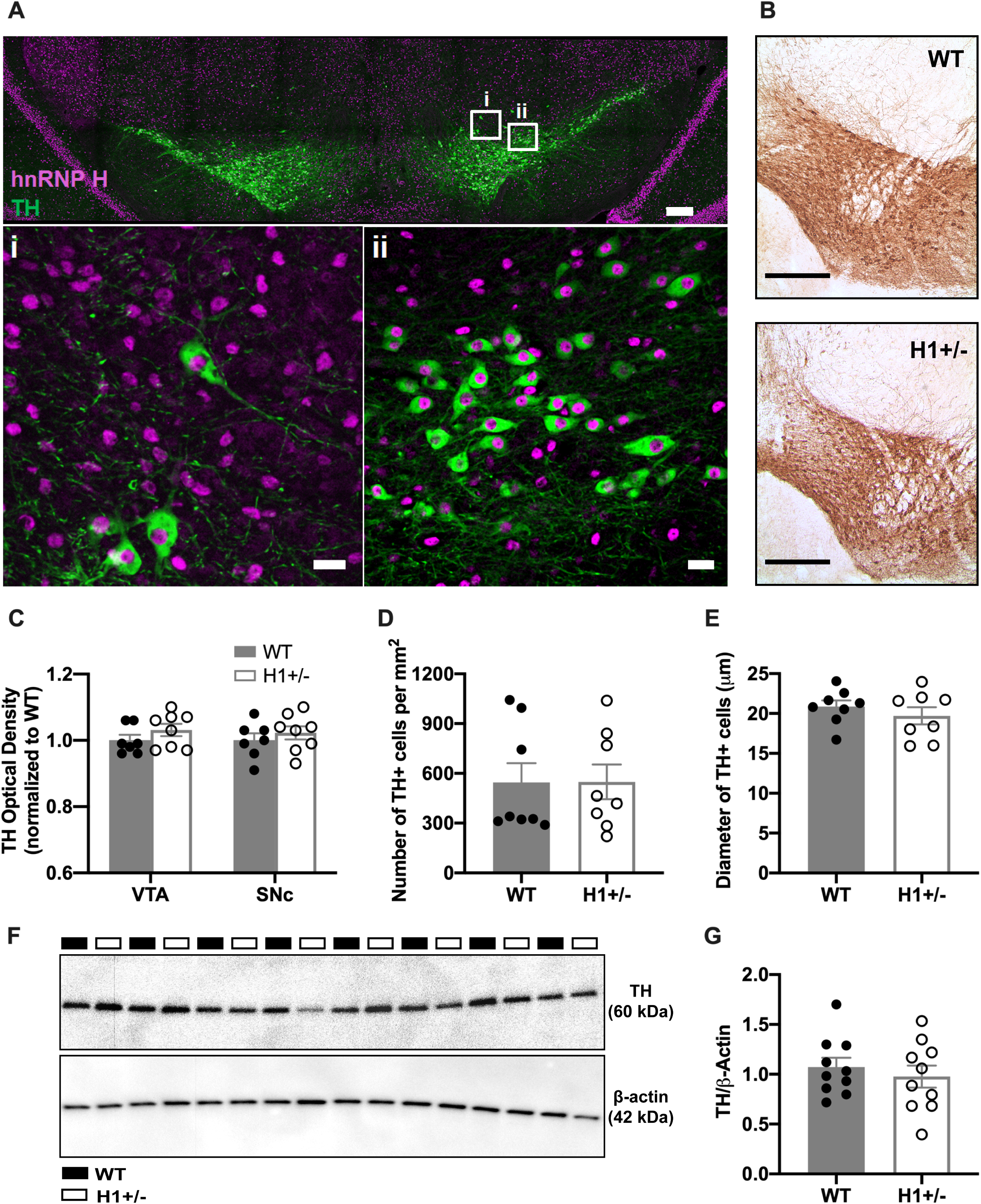
TH levels in the midbrain of H1+/- mice. No difference in TH level was detected in the VTA between H1+/- and WT mice. **(A):** Immunofluorescent staining of hnRNP H (*magenta*) and TH (*green*) was conducted in coronal midbrain sections (Bregma: −3.28 mm to −3.64 mm) containing the VTA dopaminergic neurons in adult H1+/- and WT mice. Higher magnification images in panels (***i***) and (***ii***) demonstrate nuclear expression of hnRNP H across all TH-positive dopaminergic neurons that we examined. Scale bars represent 200 μM (*top*) and 20 μM (*bottom)*. **(B):** Representative image showing immunohistochemical DAB staining of TH in coronal sections of the midbrain region (Bregma: −3.28 mm to −3.64 mm). Scale bars represent represent 1 mm. **(C-E):** Immunohistochemical DAB staining of TH in the midbrain regions revealed no genotypic difference in TH optical density **(C)**, in number of TH-positive cells **(D)**, or in the diameter of TH-positive cells **(E)** between the H1+/- and WT mice [WT, n = 8; H1+/-, n = 8; t(14) < 1, all p’s > 0.05; unpaired Student’s t-test]. **(F-G):** Immunoblot for TH protein level in the midbrain. Representative immunoblot for TH shown in **(F)** with quantification shown in **(G)**. There was no change in TH protein level in the midbrain region of H1+/- relative to WT mice [WT, n = 10; H1+/-, n = 10; t(18) = 0.67, p = 0.51; unpaired Student’s T-test].

**Figure 6.**
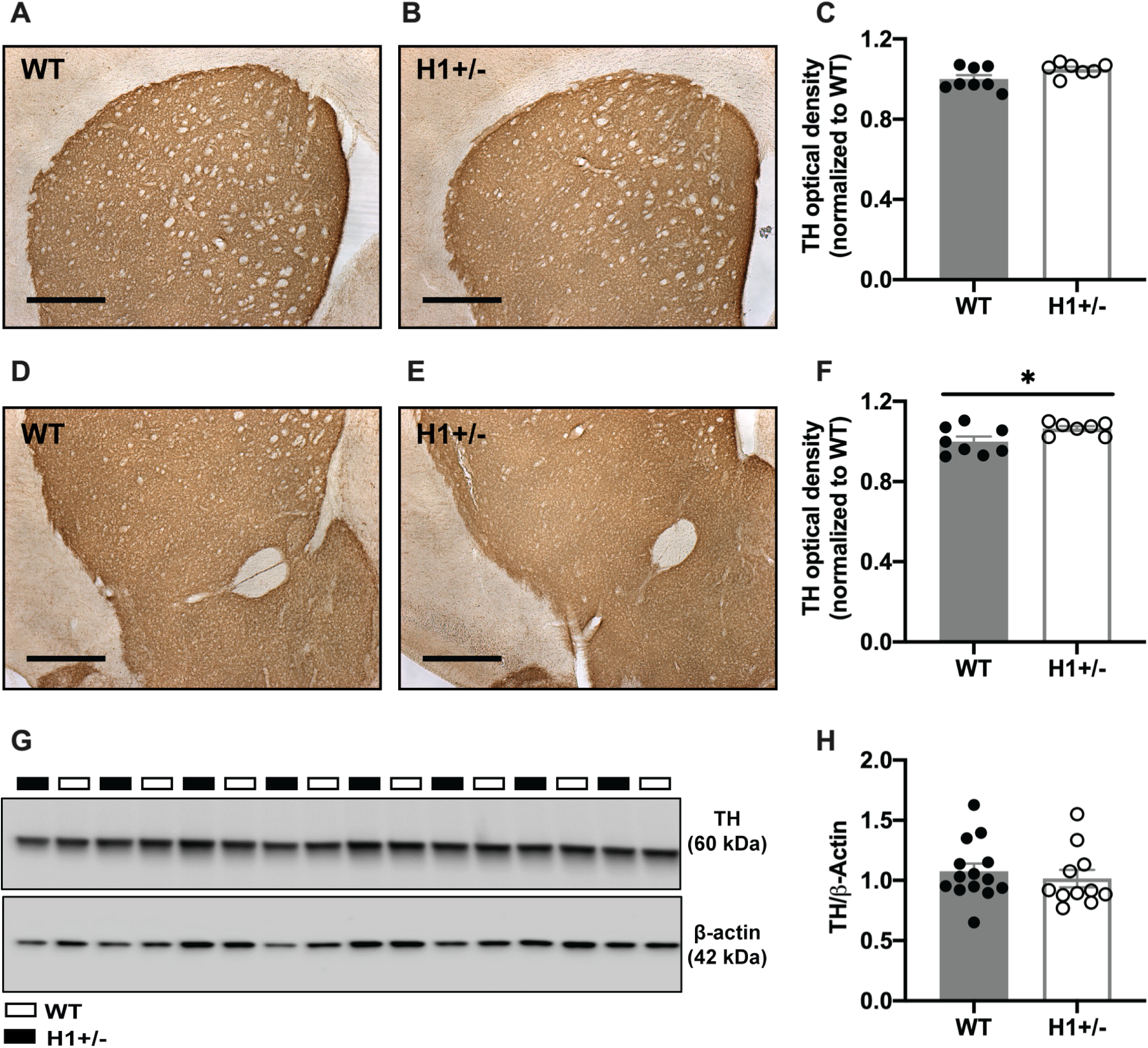
TH levels in striatum of H1+/- mice. No difference in TH level was detected in the striatum between the H1+/- versus WT mice. **(A-F):** Representative images showing immunohistochemical DAB staining on coronal sections of striatum (Bregma: 1.18 mm to 0.86 mm). **A-C:** dorsal striatum. **D-F:** ventral striatum which includes NAc. Scale bars represent represent 1 mm. Optical density (OD) analysis revealed a nonsignificant increase in TH OD in the dorsal striatum [**C:** WT, n = 7; H1+/-, n = 8; t(13) = 2.07, p = 0.10; unpaired Student’s T-test] and a small but significant increase in TH intensity in the ventral striatum of H1+/- compared to WT mice [**F**: WT, n = 7; H1+/-, n = 8; t(13) = 2.30, *p = 0.04; unpaired Student’s T-test]. **(G-H):** Immunoblot for TH protein level in the striatum. Representative immunoblot for TH shown in **(G)** with quantification shown in **(H)**. There was no change in TH protein expression in the striatum of H1+/- compared to WT mice [WT, n = 14; H1+/-, n = 11; t(23) = 0.62, p = 0.54; unpaired Student’s T-test].

An important message gleaned from the above results is that the behavioral and neurochemical deficits exhibited by H1+/- mice appear to manifest only under the influence of MA. In further support of this notion, a screen of WT and H1+/- mice in a behavioral test battery did not detect any genotypic differences in sensorimotor-gating, anxiety-like behaviors, depressive-like behaviors, or in sensorimotor coordination (**Figure 3-3**). The null results from this behavioral battery, combined with the lack of genotypic differences in saline-induced locomotion/response to a novel environment, and in various indices of basal DA transmission in the striatum argue for an active, MA-induced cell biological mechanism linking *Hnrnph1* function to MA behavior.

### Methamphetamine-induced changes in total and synaptic level of hnRNP H

The above neuroanatomical studies failed to support a neuroanatomical hypothesis of reduced neurodevelopment of DA projection pathways that could underlie reduced MA-induced DA release and behavior in H1+/- mice. Thus, to further explore the possibility of a synaptic, MA-induced cell biological mechanism that could underlie H1+/- behavior, we next examined changes in the synaptic localization of hnRNP H which is potentially relevant for understanding how H1+/- could alter MA-induced DA release. Surprisingly, we identified a two-fold *increase* in hnRNP H protein level in the striatal synaptosome of H1+/- mice, regardless of treatment (Figure 7 A-B). In contrast, there was no significant genotypic difference in hnRNP H protein level in bulk striatal tissue in response to saline or MA (Figure 7 C-D), indicating a change in localization rather than total hnRNP H protein in H1+/- mice.

**Figure 7.**
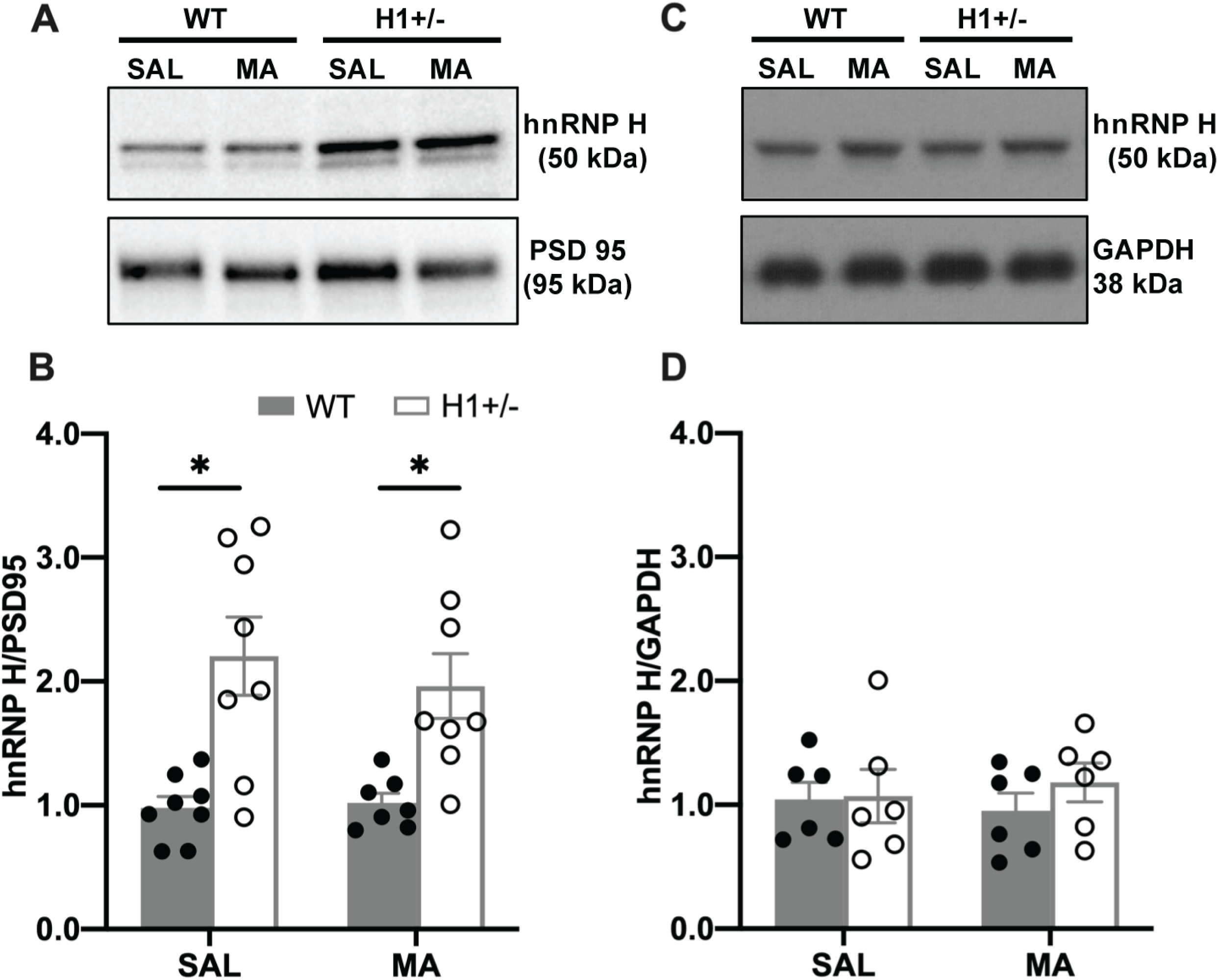
MA-induced changes in hnRNP H protein expression in striatal tissue and striatal synaptosomes of H1+/- mice. An increase in hnRNP H protein level was detected in the striatal synaptosome of H1+/- versus WT mice but no change in hnRNP H protein from total striatal tissue. **(A-B):** Representative immunoblot for hnRNP H protein level in the striatal synaptosome 30-min post saline (SAL) or MA treatment shown in **(A)** with quantification shown in **(B)**. Genotypic difference in hnRNP H protein level was detected in the striatal synaptosome of H1+/- and WT mice [main effect of Genotype: F(1,27) = 24.36, p < 0.001; main effect of Treatment: F(1,27) = 0.23; Genotype x Treatment: F(1,27) = 0.41, p = 0.53]. Collapsing across Treatment, an increase in hnRNP H protein was noted in the striatal synaptosome of H1+/- versus WT mice regardless of Treatment [t(29) = −5.06, *p < 0.001, unpaired Student’s T-test]. This finding was subsequently validated in multiple replications. **(C-D):** Representative immunoblot for *total* hnRNP H protein level the striatum 30-min post-SAL or post-MA shown in **(C):** with quantification shown in **(D).** There was no change in total striatal level of hnRNP H relative to WT mice after SAL or MA treatment [Genotype x Treatment: interaction [F(1,20) = 0.37, p = 0.55; main effect of Genotype: F(1,20) = 0.60, main effect of Treatment: F(1,20) = 0.01, p = 0.95].

To identify changes in the levels of other synaptic proteins that could mechanistically link the robust increase in synaptic hnRNP H with decreased MA-induced DA release and behavior, we examined genotypic differences in the synaptosomal proteome in the striatum of H1+/- and WT mice treated with MA versus saline using LC-MS/MS. At the behavioral level, H1+/- showed reduced MA-induced locomotor activity only in response to MA but not to saline (**Figure 8-1**). Overall, proteomic analysis of the main effect of Genotype identified a highly enriched upregulation of mitochondrial proteins in the H1+/- striatal synaptosome regardless of treatment (Figure 8A). Enrichment analysis for the set of top differentially expressed proteins (absolute log_2_FC > 0.2; p < 0.05) between H1+/- and WT mice revealed a highly significant enrichment for mitochondrial respiratory chain complex I assembly (Table 1). In examining MA-induced changes in synaptic proteins in H1+/- versus WT mice, we again identified an enrichment for alterations in metabolic processes involving components of mitochondrial complex I and V (Figures 8B**, 8-3, and** Table 2). Interestingly, in response to MA, there was a decrease in the level of mitochondrial proteins in the H1+/- mice, but an increase in the WT mice (Figure 9). The findings were independently validated with a separate cohort of mice, which also pointed to a trending, though non-significant pattern of an increase in expression of all three mitochondrial proteins that we tested in saline-treated H1+/- mice and a trending decrease in all three mitochondrial proteins in response to MA (**Figure 9-1**).

**Table 1.**
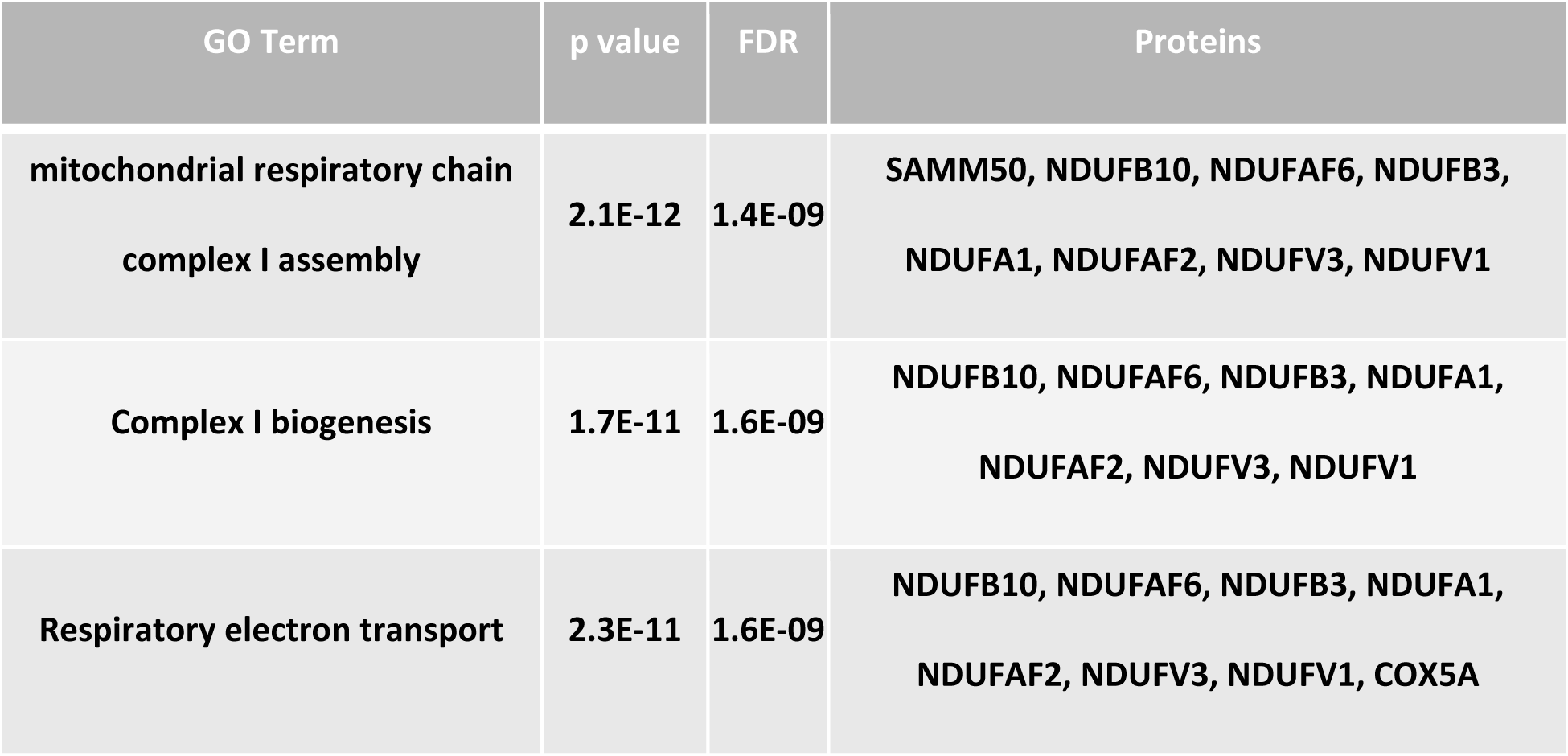
Differentially expressed proteins in H1+/- versus WT. Table showing the top three gene ontology (GO) terms for the top differential expressed protein (shown in Figure 8A) in the H1+/- versus WT striatal synaptosome.

**Table 2.**
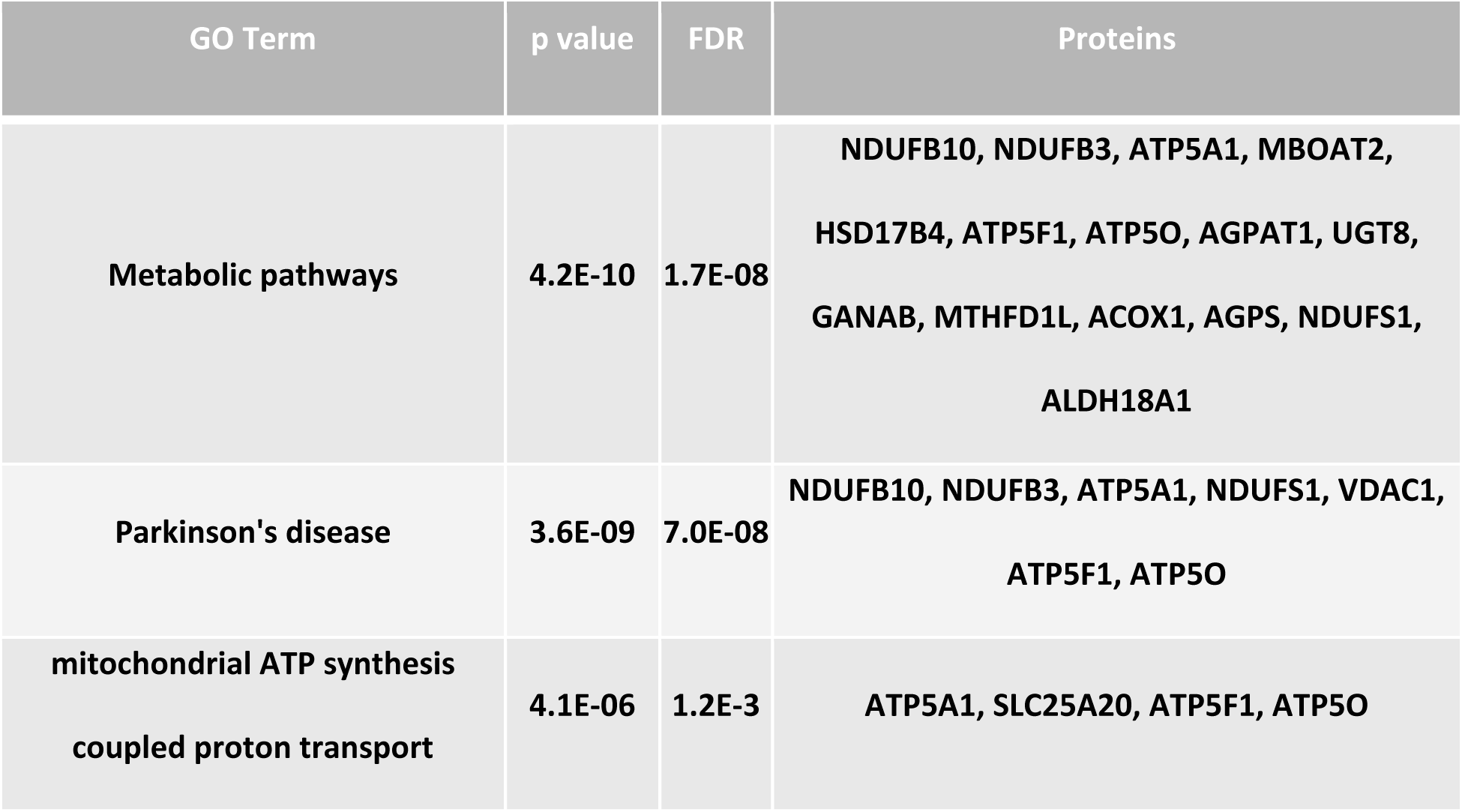
Differentially expressed proteins in [H1+/- _(MA)_ – H1+/- _(SAL)_] – [WT_(MA)_ – WT_(SAL)_]. Table showing the top 3 GO terms for the top differential expressed proteins (shown in Figure 8B) in the H1+/- versus WT striatal synaptosome in response to MA.

| GO Term | p value | FDR | Proteins |
| --- | --- | --- | --- |
| Metabolic pathways | 4.2E-10 | 1.7E-08 | NDUFB10, NDUFB3, ATP5A1, MBOAT2,<br>HSD17B4, ATP5F1, ATP5O, AGPAT1, UGT8,<br>GANAB, MTHFD1L, ACOX1, AGPS, NDUFS1,<br>ALDH18A1 |
| Parkinson's disease | 3.6E-09 | 7.0E-08 | NDUFB10, NDUFB3, ATP5A1, NDUFS1, VDAC1,<br>ATP5F1, ATP5O |
| mitochondrial ATP synthesis<br>coupled proton transport | 4.1E-06 | 1.2E-3 | ATP5A1, SLC25A20, ATP5F1, ATP5O |

**Figure 8.**
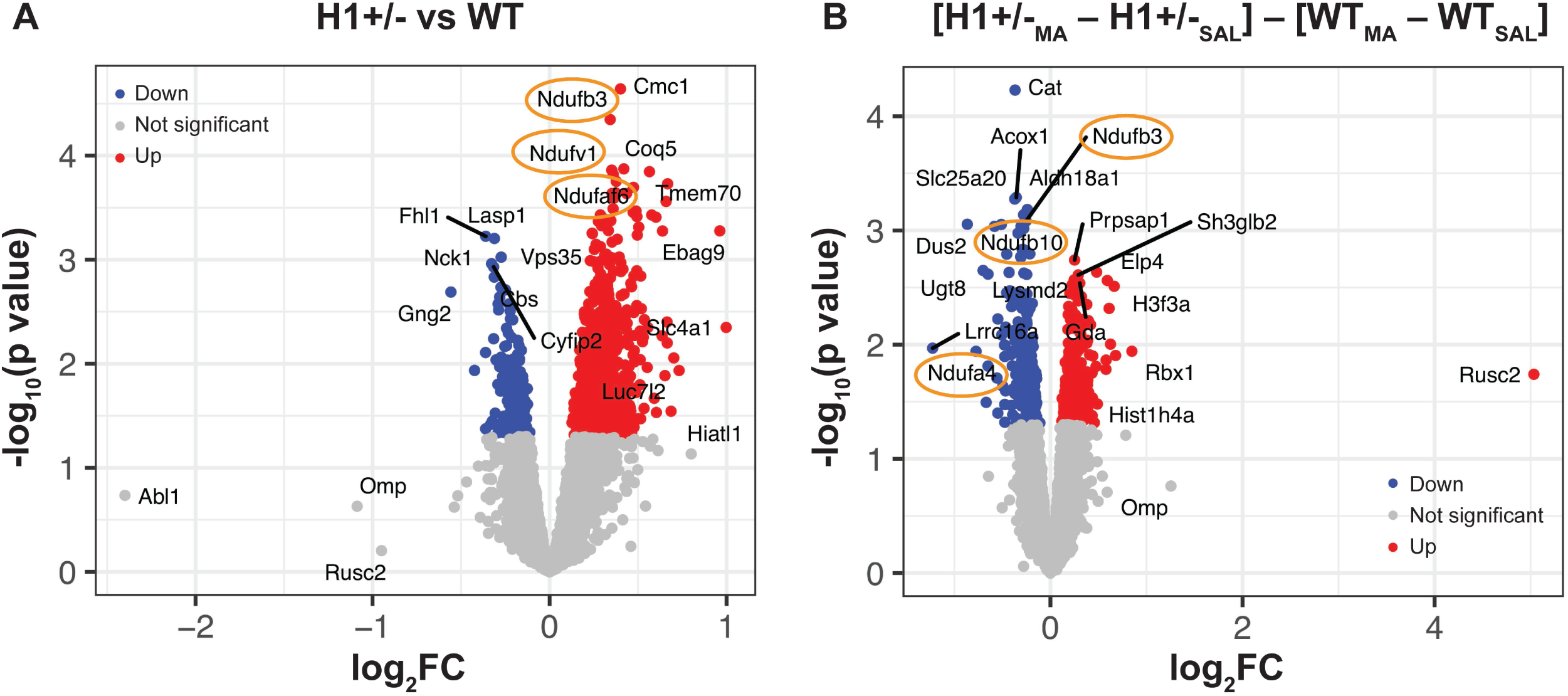
Proteomic analysis of the striatal synaptosome in H1+/- mice. On Days 1 and 2, mice were injected (i.p.) with saline (SAL) and placed into apparatus for 1 h. On Day 3, mice were injected (i.p.) with 2 mg/kg MA and placed into the apparatus for 30 min and then whole striatum was rapidly harvested via free form dissection. Locomotor activity for all three days is shown in Figure S12. The LIMMA package was used for differential analysis with Genotype and Treatment as factors. A ranked list was generated from the analysis and the fgsea R package was used to perform pre-ranked analysis, with proteins filtered for absolute log_2_FC > 0.2 and p < 0.05. **(A):** Proteomic analysis was performed to identify differences in protein abundance in the striatal synaptosome of H1+/- versus WT mice. The volcano plot shows the top differentially expressed proteins in the H1+/- versus WT striatal synaptosome. Proteins that are part of the mitochondrial respiratory complex I are circled in orange. **(B):** Proteomic analysis was performed to examine Genotype x Treatment interactions in protein abundance in the striatal synaptosome of H1+/- versus WT mice. This analysis accounted for baseline differences by examining the difference of the difference between the H1+/- and WT in response to MA: (H1+/- _MA_ – H1+/-_SAL_) – (WT_MA_ – WT_SAL_). Volcano plot showing the top differential expressed proteins between the H1+/- versus WT striatal synaptosome in response MA. Proteins that are part of the mitochondrial respiratory complex I are circled in orange.

**Figure 9.**
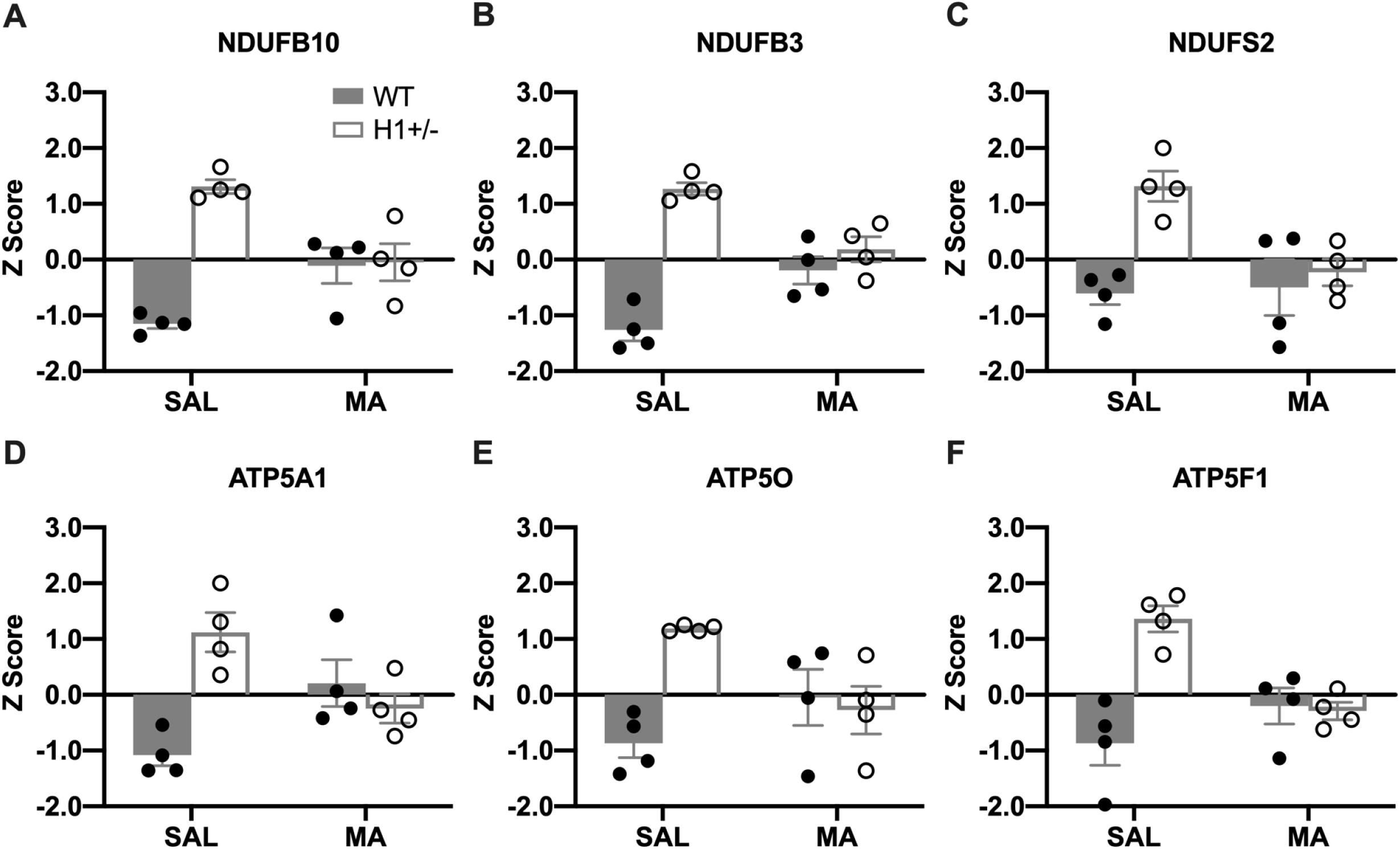
Protein expression profiles of select mitochondrial proteins in the striatal synaptosome of H1+/- and WT mice from the proteomic dataset. The protein abundance for the mitochondrial proteins are shown for the four groups: WT (SAL), H1+/- (SAL), WT (MA) and H1+/- (MA). Opposing Genotype x Treatment effects on protein levels are shown for the mitochondrial complex I components **(A-C)** and complex V ATPase subunits **(D-F)**. MA induced a decrease in protein expression of these subunits in H1+/- mice and an increase in WT mice.

## DISCUSSION

Here, we extend a role for *Hnrnph1* in MA reinforcement and reward (Figure 10). A heterozygous frameshift 16 bp deletion of the first coding exon of *Hnrnph1* reduced MA oral operant self-administration and MA-induced CPP (Figures 1 and 2) and induced a robust reduction in MA-induced DA release in the NAc (Figure 3). This DA anomaly occurred without any differences in 1) basal extracellular DA (Figure 3A-B); 2) basal DA content within striatal tissue (Figure 4A); 3) DA uptake (Figure 4B); 4) the number or staining of midbrain DA neurons within the mesolimbic and nigrostriatal dopaminergic circuits (Figure 5) or 6) any robust changes in the number of forebrain striatal puncta originating from these neurons (Figure 6). The combined results suggested an alternate mechanism underlying the decreases in MA- induced DA release and behavior in H1+/- mice. In further support of a MA-induced cell biological mechanism, there was no effect of H1+/- on spontaneous locomotion, anxiety- and depressive-like behaviors, or sensorimotor function (**Figure 3-3**).

**Figure 10.**
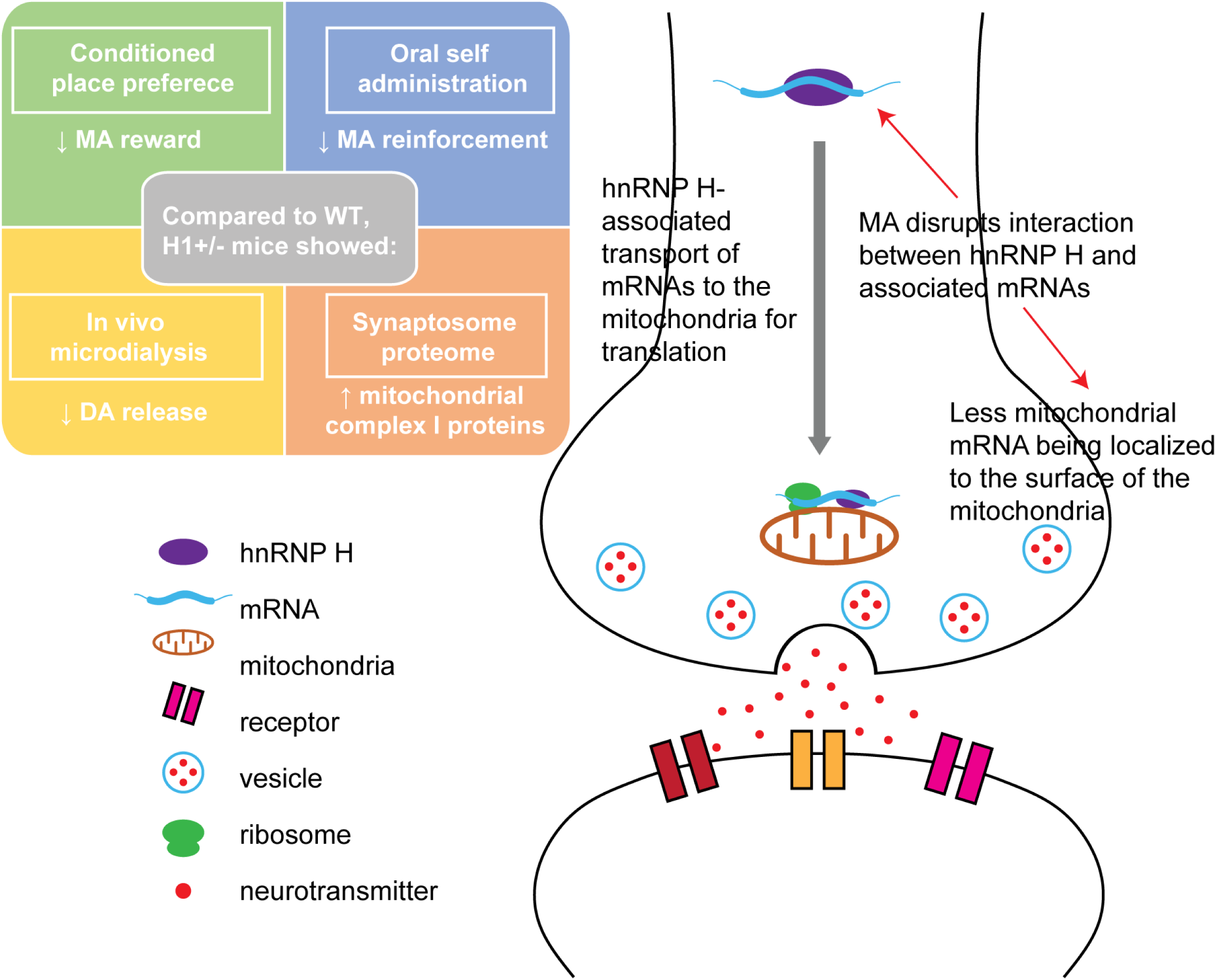
Proposed model linking increased synaptic mitochondria with a decrease in MA-induced DA release and motivated behaviors. Compared to WT mice, H1+/- mice showed reduced MA-induced DA release in the NAc and reduced sensitivity to the locomotor stimulant, rewarding, and reinforcing effects of MA. In addition, H1+/- mice show an increase in hnRNP H protein in the striatal synaptosome, with no change in total hnRNP H. This increased localization of synaptosome hnRNP H is associated with an increase in complex I mitochondrial proteins. RBPs such as hnRNP H chaperone mRNAs to membrane-bound organelles such as the mitochondria for translation and subsequent assembly in the mitochondria. In the absence of MA, hnRNP H binds to mRNA transcripts encoding mitochondria proteins and transports them to the surface of the mitochondria. MA administration disrupts hnRNP H-RNA interactions which results in less mitochondrial mRNAs being transported to the mitochondria for translation.

H1+/- mice showed less MA-CPP at 0.5 mg/kg but greater CPP at 2 mg/kg than WT mice (Figure 2). One interpretation is that MA-CPP exhibits an inverted U-shaped dose-response curve (Uhl et al., 2014) in WT and that H1+/- shifts this curve to the right, yielding reduced sensitivity to positive and negative motivational valence of MA at lower versus higher doses. Consistent with this interpretation, H1+/- mice exhibited blunted operant oral MA reinforcement. The combined data are consistent with H1+/- mice self-administering less MA because they are less sensitive to the physiological and interoceptive effects of MA due to a reduced MA-induced DA release.

H1+/- mice exhibited a blunted DA response to 0.5 and 2 mg/kg MA (Figure 3A-B). This effect on MA-induced DA release could not be explained by alterations in total DA levels (Figure 4A), or in DAT levels or function at the presynaptic membrane (Figures 4B-D **and 4-1**). An alternative explanation to DAT dysfunction is that the *Hnrnph1* mutation somehow decreases MA binding to DAT, limiting its entry into presynaptic dopaminergic neuronal terminals and decreasing DA release. Future studies are necessary to examine MA binding to DAT in H1+/- mice.

Our findings support a dopaminergic mechanism underlying reduced MA reward and reinforcement in H1+/- mice. Nevertheless, we acknowledge the potential involvement of additional neurotransmitter systems and brain regions. We did not identify any difference in basal or MA-induced changes in extracellular glutamate levels in the NAc of H1+/- mice. While MA reward is generally attributed to an increase in DA release in the NAc (Segal and Kuczenski, 1997; Adinoff, 2004), MA also increases release of norepinephrine and serotonin by targeting their respective transporters that could modulate the locomotor stimulant and/or rewarding response to psychostimulants such as MA (Haughey et al., 2002; Rothman et al., 2001; Zaniewska et al., 2015). Future studies are warranted to address these other neurotransmitters in MA-induced behavioral dysfunction in H1+/- mice as well as the possibility that H1+/- has a pleiotropic influence on behavioral (e.g., cognitive, antidepressant) and neurochemical responses to drugs targeting other membrane transporters such as NET and SERT. For example, phosphorylation of the RBP hnRNP K increases expression of SERT protein via changes in binding to the distal polyadenylation element of the transporter (Yoon et al., 2013).

The two-fold increase in hnRNP H protein in the striatal synaptosome of H1+/- mice with no change in total hnRNP H protein was surprising. This finding was observed in multiple replication studies and suggests a redistribution of hnRNP H protein to the synapse in H1+/- mice. We performed LC-MS/MS analyses on striatal synaptosomes following 2 mg/kg MA (i.p.) to further understand the effect of increased synaptosomal hnRNP H on global changes in protein expression and the underlying cell biological mechanisms. We identified a higher abundance of several mitochondrial proteins, in particular, complex I of the mitochondrial respiratory chain in H1+/- mice. The mammalian complex I consists of 38 nuclear DNA-encoded subunits (Sharma et al., 2009) and our proteomic analysis identified 8 out of the 38 subunits that showed higher expression in H1+/- mice (Figures 8-9). Proteomics has been widely used to study the effects of MA on protein expression in the brain tissues of animals (Liao et al., 2005; Faure et al., 2009; Bosch et al., 2015). Iwazaki et al. (2006) used two-dimensional gel electrophoresis proteomics and found that a single low dose of MA (1 mg/kg) administration in rats induced differential expression of proteins involved in mitochondria/oxidative metabolism. Furthermore, chronic exposure of 1 mg/kg MA induced locomotor sensitization and neurotoxicity along with a downregulation of numerous striatal proteins indicating mitochondrial dysfunction and an oxidative response (Iwazaki et al., 2007; Chin et al., 2008).

Recent studies showed postsynaptic dendritic mitochondrial fission and fusion processes mediate cellular and behavioral plasticity, spine and synapse formation, and synaptic function (Li et al., 2004; Oettinghaus et al., 2016; Chandra et al., 2017; Divakaruni et al., 2018). Dynamin-related protein (Drp1), a GTPase involved in mitochondria fission, has been shown to regulate addiction-relevant behavior during early cocaine abstinence (Chandra et al., 2017), with inhibition of mitochondrial fission blunting cocaine-seeking and locomotor sensitization. While the results of our synaptosomal proteome dataset did not reveal any mitochondrial fission and fusion mediators, it is still possible that changes in mitochondrial proteins in the post- synaptic dendrites contribute to behavioral differences. Future studies will isolate potential pre- versus postsynaptic mechanisms.

Most mitochondrial proteins are nuclear-encoded and must be transported to mitochondria for organelle-coupled translation (Williams et al., 2014). RBPs play a critical role in targeting mRNAs to membrane-bound organelles such as mitochondria. RBPs interact with mRNAs and chaperone them toward mitochondrial outer membranes where they are translated (Gerber et al., 2004; García-Rodríguez et al., 2007; Eliyahu et al., 2010; Gehrke et al., 2015). RBPs recognize and bind mitochondria-targeting RNA elements to form higher-order units called mRNA ribonucleoprotein (RNP) complexes consisting of mRNAs and associated RBPs (Béthune et al., 2019; Rossoll and Bassell, 2019). The robust increase in hnRNP H protein in the striatal synaptosome accompanied by the increase in several complex I subunits suggests a novel function for hnRNP H in targeting mRNAs encoding for subunits of mitochondrial complex I to the mitochondria, thus, regulating local translation (Figure 10). Once the mRNAs are transported nearby the mitochondria, hnRNP H could coordinate with other RBPs to form an RNP complex to stabilize mRNAs to the mitochondrial membrane where translational activators initiate translation. An example of a RBP with such function is Puf3pm that binds to *ATP2* mRNA encoding mitochondrial components of the F_1_F_0_ ATPase and localizes to the outer mitochondrial membrane (García-Rodríguez et al., 2007). Future studies involving cross-linking immunoprecipitation combined with RNA-seq (CLIP-seq) will identify target mRNAs bound to hnRNP H and determine the degree of enrichment for mRNAs encoding mitochondrial complex I subunits.

Mitochondria are abundant at the synapse where they generate ATP for Ca^2+^ buffering, vesicle release, and recycling (Vos et al., 2010; Devine and Kittler, 2018). Mitochondria consist of five oxidative phosphorylation complexes (I through V) (Mimaki et al., 2012; Sharma et al., 2009). Complex I is the first enzyme of the respiratory chain and initiates electron transport continuing to complex II through IV to generate redox energy for Complex V to produce ATP. The increase in Complex I subunits in H1+/- mice could increase Complex I activity and synaptic ATP production to support vesicle fusion, DA release, and DA transport back into the cells via DAT and Na+/K+ ATPase to counteract MA-induced DA release. We found a Genotype by Treatment effect on protein levels of F_0_F_1_ ATP synthase subunits (Atp5a1, ATP5f1 and Atp5o) of Complex V (Figure 9) in the synaptosomal proteome whereby MA decreased these ATP synthase subunits in H1+/- mice which could decrease ATP production in response to MA in H1+/- mice and affect extracellular DA levels.

Besides binding and targeting RNAs to the mitochondria for translation, hnRNP H, like other RBPs, could bind and target proteins via its glycine-rich domain in an activity-dependent manner (Tiruchinapalli et al., 2008). In an animal model for frontotemporal dementia, ploy(GR) aggregates (resulting from hexanucleotide repeats in *C9ORF72)* bind to mitochondrial complex V protein ATP5A1 to increase its ubiquitination and degradation through the proteasome pathway, thus disrupting mitochondrial function (Choi et al., 2019). The higher level of synaptosomal hnRNP H protein in H1+/- mice could increase binding to mitochondrial complex I and V proteins (e.g., via the glycine-rich domain) to prevent degradation, yielding higher protein levels at baseline (Figures 8 and 9). MA administration could then decrease hnRNP H-protein interactions in H1+/- mice, thus decreasing synaptic mitochondrial proteins and synaptic function.

Taken together, the opposing effects of MA treatment on synaptic abundance of mitochondrial complex I proteins between H1+/- and WT mice could represent a mechanism underlying blunted MA-induced DA release in H1+/- mice. Future studies will focus on the interaction between hnRNP H and mRNA encoding mitochondrial complex I and V subunits at the protein-RNA and protein-protein level (pre- and postsynaptically) and determine whether disruption of such interactions can alter ATP production and DA release.

## Acknowledgments

This study was supported by National Institutes of Health/National Institute on Drug Abuse (NIH/NIDA) Grant Nos. R00DA029635 (C.D.B.), R01DA039168 (C.D.B.), F31DA40324 (N.Y.), and N01DA-14-7788; NIH/National Institute of General Medical Sciences (NIGMS) for Grant No. T32GM008541 (N.Y., Q.T.R. and J.A.B.); and the Burroughs Wellcome Fund Transformative Training Program in Addiction Science Grant No. 1011479 (N.Y., Q.T.R. J.A.B,). We would like to acknowledge Dr. Haley Melikian for providing guidance on the dopamine transporter immunoblot. We would also like to acknowledge Drs. David Moody, Wenfang B Fang, Olga Averin, and Jonathan Crites for providing the analytical service for measuring methamphetamine concentration in our biological samples.

## EXTENDED DATA LEGENDS

**Figure 1-1. Bitter/Quinine taste sensitivity in H1+/- mice.** There were no genotypic differences between H1+/- and WT mice in average quinine intake (0.003-0.6 mg/ml) [Quinine effect: F(5,100) = 44.63, p < 0.0001; Genotype effect: F(1,20) = 2.07, p = 0.17; Genotype x Quinine interaction: F(5,100) = 0.50, p=0.78].

**Figure 1-2. High-dose MA oral self-administration in H1+/- mice.** Acquisition of oral MA self-administration with initial training to respond for 200 mg/mL. Mice were trained to nose-poke under an FR1 schedule of reinforcement (20 s time-out) for 20 uL delivery of a 200 mg/L MA solution over the course of 2 weeks. **(A and B):** While both MA intake and the relative amount of reinforced responding varied across days [for intake, Day effect: F(13,273)=8.39, p<0.0001; for % reinforced responding, F(13,273)=16.20, p<0.0001], no genotypic differences were detected for either variable during this phase of testing [no Genotype effects or interactions, p’s>0.50]. Furthermore, there was no difference between the number of WT (2 of 12) versus H1+/- (3 of 11) mice that failed to meet our acquisition criteria [>10 responses with 70% of responding directed at the MA-reinforced hole; λ2=0.24, p=0.62].

**Figure 2-1. Locomotor activity in the CPP box on Day 1, 8, and 9 for 0 mg/kg (saline, SAL), 0.5 mg/kg MA, and 2 mg/kg MA in H1+/- mice.** For all data shown, mice were allowed open access to both sides of the apparatus via an open entryway in the middle of the dividing wall. On Day 1, mice were injected with SAL and assessed for initial preference for the MA-paired side. On Day 8, mice were injected with SAL and tested for drug-free MA-CPP. On Day 9, mice were tested with either SAL or with the same dose of MA that they received during training to test for state-dependent MA-CPP. In examining the main effect of Genotype and interaction between Genotype and Time for each dose, a mixed-model two-way ANOVA with Genotype (between-subjects) and Time (repeated measure) as factors was used. **(A-C):** Locomotor activity for Day 1. No genotypic difference was detected for Day 1 locomotor activity for 0.5 and 2 mg/kg MA [SAL: F(1,45) = 24.76, p = 3.81E-5; 0.5 mg/kg MA: F(1,43) = 1.894, p = 0.176; 2 mg/kg: F(1, 33) = 2.657, p = 0.113]. **(D-F):** Locomotor activity for Day 8. Genotypic difference was detected for Day 8 locomotor activity for 0.5 mg/kg MA but not for SAL and 2 mg/kg MA [SAL: F(1,45) = 2.22, p = 0.143; 0.5 mg/kg MA: F(1,43) = 6.117, p = 0.017; 2 mg/kg: F(1, 33) = 2.897, p = 0.1]. **(G-I):** Locomotor activity for Day 9. No genotypic difference was detected for Day 9 locomotor activity for the SAL and 0.5 mg/kg MA dose [SAL: F(1,44) = 2.956, p = 0.1; 0.5 mg/kg MA: F(1,42) = 0.31, p = 0.58]. In response to 2 mg/kg MA, H1+/- showed a trend toward less total distance over the 1 h period compared to WT [main effect of Genotype: F(1,33) = 3.032, p = 0.091]. Sample sizes are indicated in the parentheses.

**Figure 3-1. DA no net flux, baseline DA levels and probe placements in H1+/- mice. (A):** DA no net flux in H1+/- and WT mice revealed no genotypic differences at the point of no DA- flux within the NAc [Genotype by Sex ANOVA, all p’s > 0.35; y = 0: 5.00 ± 0.44 for WT (n = 16) versus 4.68 ± 0.70 for H1+/- (n = 16)]. Although we failed to detect a genotypic difference in the extraction fraction (slope; Genotype and interaction, p’s>0.10], we detected a main effect of Sex [F(1,31) = 12.09, p = 0.002] that reflected a greater extraction fraction in females (0.94 ± 0.02) versus males (0.69 ± 0.04). These data do not support an effect of *Hnrnph1* deletion on basal extracellular DA content (y = 0) or basal DA release/reuptake (extraction fraction) associated with the reduced MA-induced DA release observed in H1+/- mice. **(B):** Summary of the average extracellular DA content (nM) prior to an acute injection of MA in WT and H1+/- mice. When assessed using conventional *in vivo* microdialysis procedures, we failed to detect any Genotype or Sex differences in the average basal extracellular DA content, prior to MA injection (no Genotype or Sex effects, nor any Genotype or Sex interactions, p’s>0.40). Probe recovery was lower in the 2 mg/kg MA study than it was in the 0.5 mg/kg study [Dose effect: F(1,33) = 47.83, p < 0 .0001]. Sample sizes are indicated in parentheses. **(C):** Probe placements for WT and H1+/- mice fall primarily within the NAc shell (Bregma: 1.54 mm to 1.18 mm).

**Figure 3-2. *In vivo* microdialysis of MA-induced glutamate release in H1+/- mice. (A):** Linear regression analyses of the results of a glutamate no net-flux *in vivo* microdialysis study (0, 2.5, 5 and 10 µM glutamate) revealed no genotypic differences in the x-intercept (an estimate of basal extracellular glutamate content in µM; WT: 2.28 ± 0.56, n = 7 versus H1+/- 2.55 ± 0.18, n = 8) and no difference in the extraction fraction or slope of the regression (an index of neurotransmitter release/clearance; WT: 0.73 ± 0.07 versus H1+/-: 0.80 ± 0.04). **(B-C):** No genotypic or sex differences were detected in average baseline extracellular glutamate levels, determined under conventional *in vivo* microanalysis procedures (Genotype x Sex x Dose ANOVA, all p’s>0.09; WT=3.27±0.39 µM, n=32 versus H1+/-=3.67±0.32 µM, n=30). Data are expressed as the percent change from the average baseline value to better visualize the time-course of the glutamate response to an acute injection of **(B)** 0.5 or **(C)** 2 mg/kg MA.

**Figure 3-3. Behavioral test battery in H1+/- mice. (A):** Timeline of behavioral battery testing in drug-naïve H1+/- (n = 38) and wild-type (WT) mice. **(B):** Prepulse inhibition of acoustic startle revealed no genotypic differences in the PPI response to 75Hz [*left*; t(69) = 1.26, p = 0.21] or 95Hz [*right*; t(69) < 1]. **(C):** Novel object test revealed no genotypic differences in novel object contact number [*left*; t(69) < 1] or contact time [*right*; t(69) = 1.62, p = 0.11]. **(D):** Marble burying test revealed no genotypic differences in the amount of time (s) spent burying [t(69) < 1]. **(E):** Light/dark shuttle box test revealed no genotypic differences in the amount of time (s) spent in the light zone [t(69) = 1.48, p = 0.15]. **(F):** Porsolt swim test revealed no genotypic differences in the amount of time immobile [t(69) < 1]. **(G):** In the accelerating rotorod test, there were no genotypic differences in the ten-trial average time spent on the rotorod [t(69) < 1].

**Figure 4-1. MA-induced changes in DAT protein level in the striatal synaptosome of H1+/- mice.** There was no change in DAT protein level in the striatal synaptosome of H1+/- mice **(A):** Immunoblot showing DAT protein level the striatal synaptosome at 30-min post saline (SAL) or MA treatment. **(B):** The striatal synaptosome of H1+/- mice exhibited no change in DAT relative to WT mice after SAL or MA treatment. A two-way ANOVA (Genotype x Treatment) indicated a no effect of Genotype [F(1,20) = 0.031, p = 0.86], no effect of Treatment [F(1,20) = 0.037, p = 0.85], and no significant interaction between Genotype and Treatment [F(1,20) = 1.24, p = 0.28].

**Figure 5-1. Schematics showing Bregma positions of the brain regions in IHC studies and Western blots. (A):** VTA and SNc of midbrain. Left: schematic showing the ventral midbrain region dissected for Western blot analysis. Right: coronal brain diagrams showing highlighted VTA (yellow) and SNc (teal) for IHC. **(B)**: Coronal brain diagrams showing highlighted dorsal striatum (green) and NAc (yellow) for IHC.

**Figure 6-1. Schematics and puncta count of TH IHC staining in H1+/- mice. (A)**: A 225,000 x 225,000 µm grid was overlaid onto the images in Image J, and every third field (indicated by the black dots) was graded for total number of puncta. Regions of interest were graded by subtracting the background, setting a threshold, creating a binary image, and conducting particle analysis to count total puncta number based on roundness and total size (as indicated by threshold black/white image on bottom right). **(B):** Stereological analysis on number of TH-positive puncta revealed no genotypic difference on the number of dopaminergic terminals in the dorsal and ventral striatum of H1+/- relative to WT mice [WT, n = 8; H1+/-, n = 8; t(14) < 1].

**Figure 7-1. *Hnrnph1* whole brain mRNA and hnRNP H whole body protein expression in WT, H1+/-, and H1-/- mice.** H1-/- mice and WT littermates were generated by intercrossing H1+/- and H1+/-. Embryos were harvested at E12 for genotyping using a restriction enzyme-based assay. **(A):** A PCR amplicon capturing the deleted region was digested with BstNI. WT mice had two copies of two BstNI restriction sites, and thus, restriction digest produced three fragments (58 bp, 157 bp, and 153 bp) corresponding to two bands on the gel. H1-/- mice had two copies of a single BstNI restriction sites, and thus, restriction digest produced two fragments (153 bp and 198 bp). H1+/- mice possessed one copy of each of the two BstNI restriction sites and one copy of a single BstNI restriction site, and thus, restriction digested produced 5 fragments (58 bp, 153 bp, 153 bp, 157 bp, and 198 bp) corresponding to three bands on the gel. **(B):** There was a non-significant, gene dosage-dependent increase in the transcript level *Hnrnph1* in H1+/- and H1-/- mice [F(2,9) = 2.161, p = 0.18; one-way ANOVA]. The 1.5-increase in *Hnnph1* transcript level in H1+/- mice replicated our previously published data (Yazdani et al. 2015). The > 2-fold increase in *Hnrnph1* transcript level in H1-/- with two copies of the mutation provides further functional support for increased expression of the mutant transcript [WT versus H1-/-: t(6) = −2.51, **#**p = 0.05]. **(C):** There was no genotypic difference in *Hnrnph2* transcript level [F(2,9) = 1.33, p = 0.32; one-way ANOVA]. **(D-G):** Protein expression of hnRNP H in WT, H1+/-, and H1-/- mice. There was no genotypic difference in hnRNP H protein expression using an antibody targeting the C-terminus of hnRNP H [**D-E;** F(2,9) = 0.89, p =0.44; one-way ANOVA], nor was there a significant difference using an antibody specific for the N-term of hnRNP H [**F- G**; F(2,9) = 1.64, p = 0.25; one-way ANOVA]. n = 4 for each genotype.

**Figure 7-2. Quantification of hnRNP H1 and hnRNP H2 protein expression using peptide information from mass spec of hnRNP H immunoprecipitates from H1+/- and WT striatum.** A separate cohort of animals were used for this study. Co-immunoprecipitation was formed to pull down hnRNP H and associated proteins. Using the “Similarity” function in the Scaffold software, peptides exclusive to hnRNP H1 or hnRNP H2 were identified and quantified. **(A):** A list is shown for the peptides that are unique to hnRNP H1. **(B):** Quantification of peptides unique to hnRNP H1. There was a trend for a decrease in peptide GLPWSCSADEVQR in the striatum of H1+/- versus WT mice [WT, n = 3; H1+/-, n = 3; t(4) = 1.938, **#**p = 0.06; unpaired Student’s T-test]. Amino acids PWSCS within this peptide are encoded by the deleted region (GCCCTGGTCCTGCTCC) in exon 4 of *Hnrnph1* in the H1+/- mice. **(C):** Table outlining the peptides that are unique to hnRNP H2. **(D):** Quantification of hnRNP H2 unique peptides. No change in unique peptides of hnRNP H2 was observed between WT and H1+/- mice [WT, n = 3; H1+/-, n = 3; t(4) < 1; unpaired Student’s T-test].

**Figure 8-1. MA-induced locomotor activity in H1+/- and WT mice that were used for striatal synaptosome harvest for proteomic analysis.** On Days 1 and 2, mice were injected (i.p.) with saline (SAL) and placed into apparatus for 1 h. On Day 3, mice were injected (i.p.) with 2 mg/kg MA and placed into apparatus for 30 min followed by subsequent removal of the striatum. **(A-B):** Locomotor activity for Day 1 **(A)** and Day 2 **(B)** for 1 h in 5-min bin. No difference in total distance traveled across [no interaction between Genotype x Treatment; Day 1: F(1,12) = 0.17; Day 2: F(1,12) = 0.21]. **(C):** Locomotor activity for Day 3 for 30 min in 5-min bin. H1+/- and WT showed Genotype x Treatment difference in sensitivity to SAL and MA [main effect of Genotype: F(1,12) = 8.53, p = 0.013; main effect of Treatment: F(1,12) = 77.42, p < 0.001; Genotype x Treatment: F(1,12) = 5.85, p = 0.03]. In response to MA (2 mg/kg, i.p.), H1+/- showed less distance traveled compared to WT [main effect of Genotype: F(1,6) = 7.84, p = 0.03]. There was a significant MA-induced decrease in locomotor activity in the H1+/- mice [t(6) = −2,46, −2.45, −3.02 for 20, 25 and 30-min time point respectively, *p < 0.05]. However, in response to SAL, there was no genotypic difference in locomotor activity [main effect of Genotype: F(1,6) = 0.70], all p’s > 0.05].

**Figure 8-2. Concentration of MA and amphetamine in the striatum at 30 min post-MA injection in H1+/- mice.** On Days 1 and 2, mice were injected (i.p.) with saline and placed into apparatus for 1 h. On Day 3, mice were injected (i.p.) with 2 mg/kg MA and placed into apparatus for 30 min followed by subsequent removal of the striatum. **(A):** No genotypic difference was detected in the striatal concentration of MA [t(14) = 0.72, p = 0.48; unpaired Student’s T-test]. **(B):** Also, no genotypic difference was detected in the striatal concentration of amphetamine [t(14) = 0.22, p = 0.83; unpaired Student’s T-test].

**Figure 8-3. Network and pathway analysis of synaptosomal proteome of H1+/- versus WT mice treated with MA or SAL [(H1+/-_MA_ – H1+/-_SAL_) – (WT_MA_ – WT_SAL_)].** Enrichment results for the comparisons were visualized using Cytoscape and EnrichmentMap. Pathways were clustered and annotated with themes using AutoAnnotate. Nodes (grey circle) represent the pathways and the size of each node represents the number of proteins. Each circle within each node represents the individual protein (red is up and blue is down relative to WT). An enrichment for metabolic processes is detected in the H1+/- mice and WT in response to MA in which MA decreases the expression of proteins involved in metabolic processes in the H1+/- mice. The cutoff for the differential analysis is set to absolute log_2_FC = 0.2 and p < 0.05.

**Figure 9-1. Immunoblots of select mitochondrial proteins in the synaptosome of H1+/- and WT mice.** Three mitochondrial proteins (NDUFS2, ATP5A, a d ATP5F1) were selected for independent validation of mass spectrometry results in a separate sample cohort via immunoblot. **(A):** Immunoblots for NDUFS2, ATP5A1 and ATP5F1. **(B-D):** Quantification of protein expression of three select mitochondrial proteins. All three proteins show the same trend as the mass spectrometry results depicted in Figure 9 where an increase in expression was detected in the SAL-treated H1+/- mice. **(B):** NDUFS2: t(6) = −1.39 p = 0.11; **(C)** ATP5A1: t(6) = −1.64, p = 0.08; **(D)** ATP5F1: t(6) = −1.24, p = 0.13]. n = 4 per genotype per dose.

