## Supplement Figures for "A mutation in *Hnrnph1* that decreases methamphetamine-induced reinforcement, reward, and dopamine release and increases synaptosomal hnRNP H and mitochondrial proteins" (from another version of this preprint: version 1, 2019-07-28)

### Figure S1

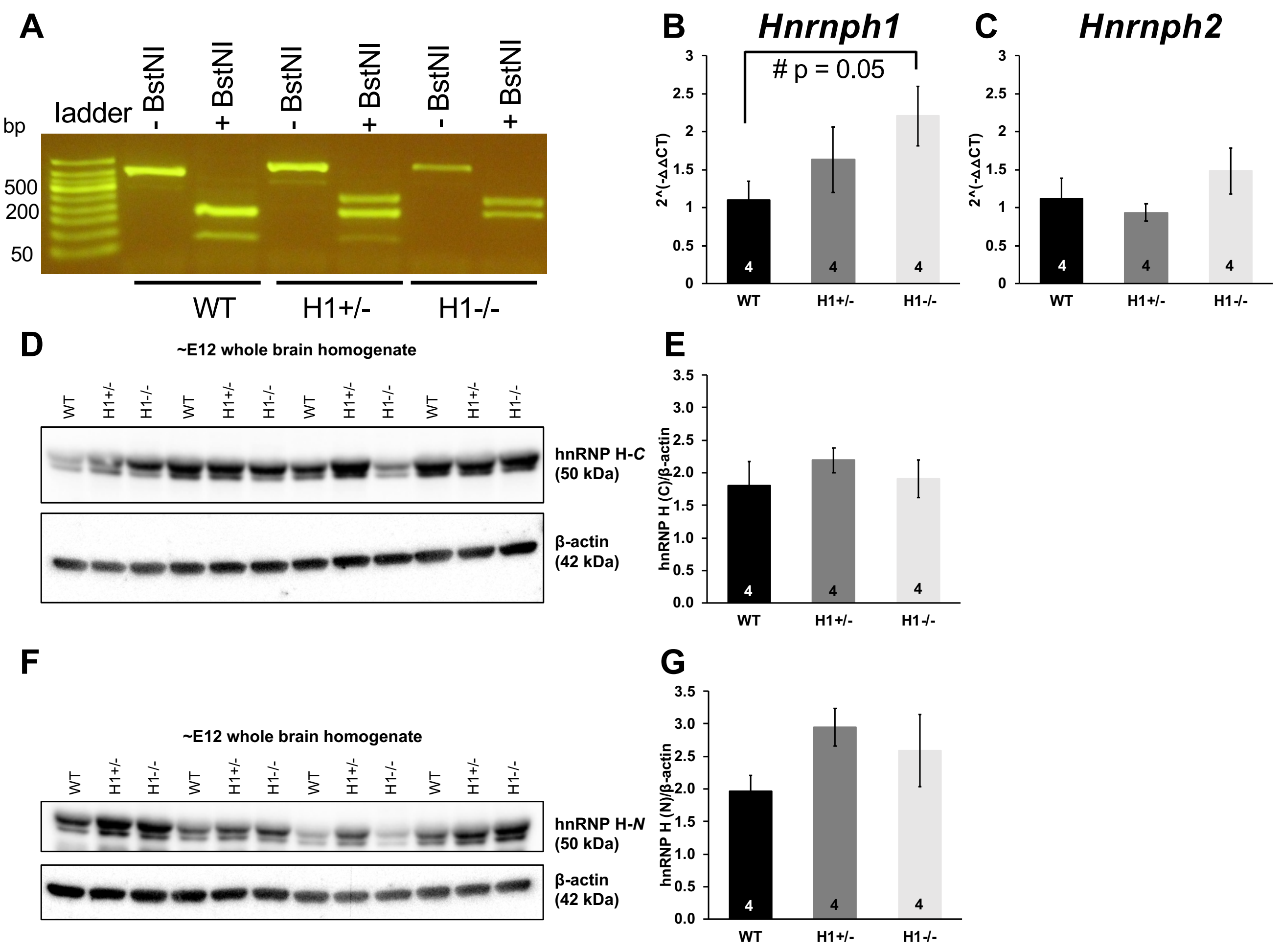
A
B
C
D
E
F
G
ladder
- BstNI
+ BstNI
bp
500
200
50
WT
H1+/-
H1-/-
Hnrnph1
Hnrnph2
# p = 0.05
2^(-ΔΔCT)
~E12 whole brain homogenate
hnRNP H-C
(50 kDa)
β-actin
(42 kDa)
hnRNP H (C)/β-actin
hnRNP H-N
(50 kDa)
hnRNP H (N)/β-actin

### Figure S2

A

| | hnRNP H1 unique peptide sequence |
|---|---|
| 1 | MMLGAEGGEGFVVK |
| 2 | MLGAEGGEGFVVK |
| 3 | GL**PWSCS**ADEVQR |
| 4 | SNNVEMDWVLK |
| 5 | GAYGGGYGGYDDYNGYNDGYGFGSDR |
| 6 | YGDGGSTFQSTTGHCVHMR |
| 7 | VTGEADVEFATHEDAVAAMSK |
| 8 | YVELFLNSTAGASGGAYEHR |

B

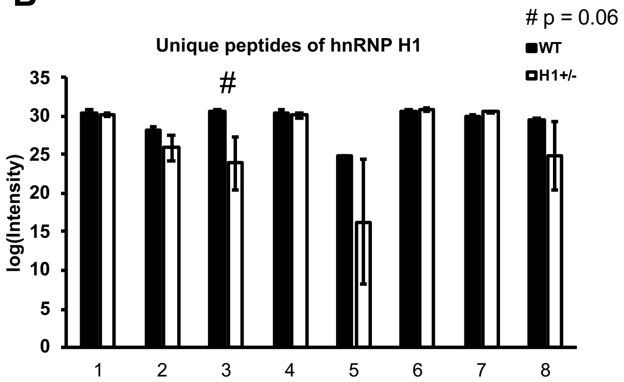


C

| | hnRNP H2 unique peptide sequence |
|---|---|
| 1 | ATENDIYNFFSPLNPMR |
| 2 | GAYGGGYGGYDDYGGYNDGYGFGSDR |
| 3 | GLPWSCSAEEVMR |
| 4 | SNSVEMDWVLK |
| 5 | VTGEADVEFATHEDAVAAMAK |
| 6 | YGDGGSSFQSTTGHCVHMR |

D

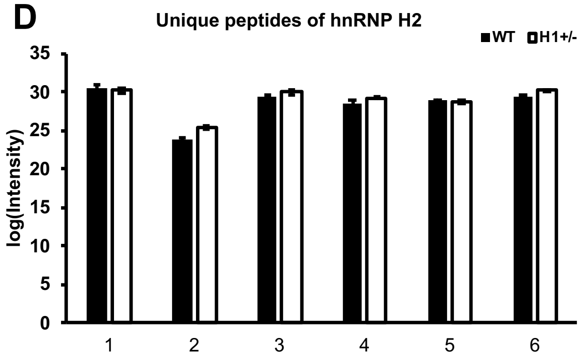

### Figure S3

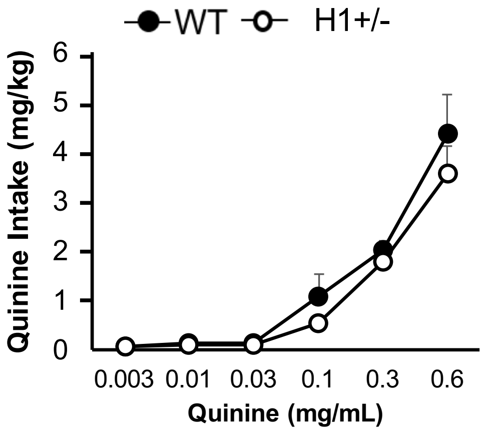
WT
H1+/-
Quinine Intake (mg/kg)
0
1
2
3
4
5
6
0.003
0.01
0.03
0.1
0.3
0.6
Quinine (mg/mL)

### Figure S4

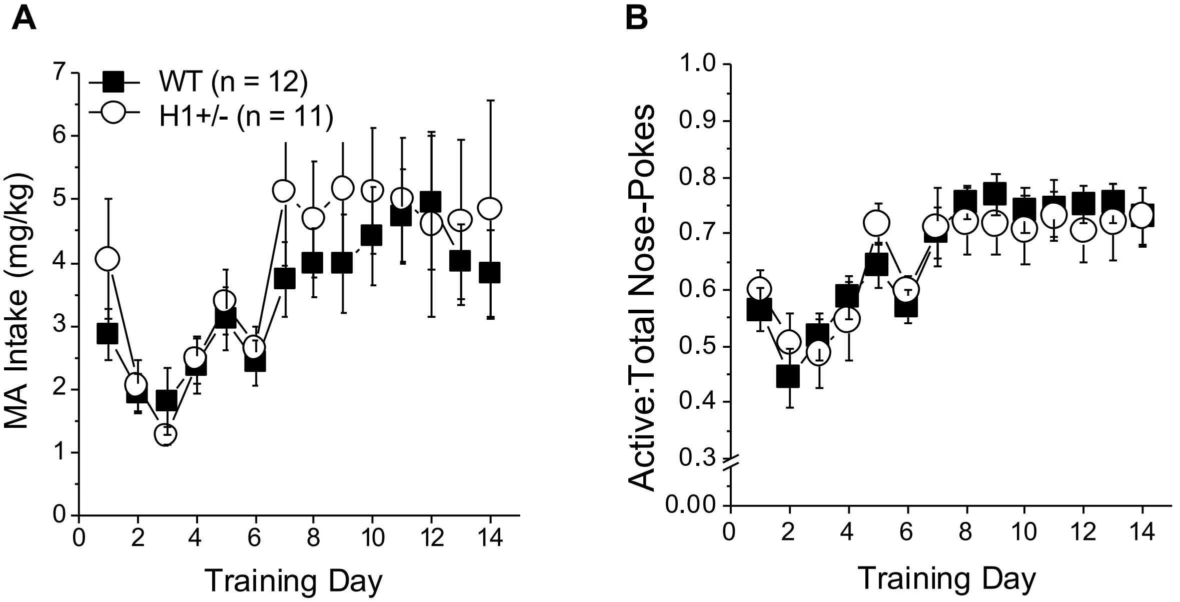
A
B
WT (n = 12)
H1+/- (n = 11)
MA Intake (mg/kg)
Active:Total Nose-Pokes
Training Day
Training Day
0
1
2
3
4
5
6
7
0.00
0.3
0.4
0.5
0.6
0.7
0.8
0.9
1.0
0
2
4
6
8
10
12
14

### Figure S5

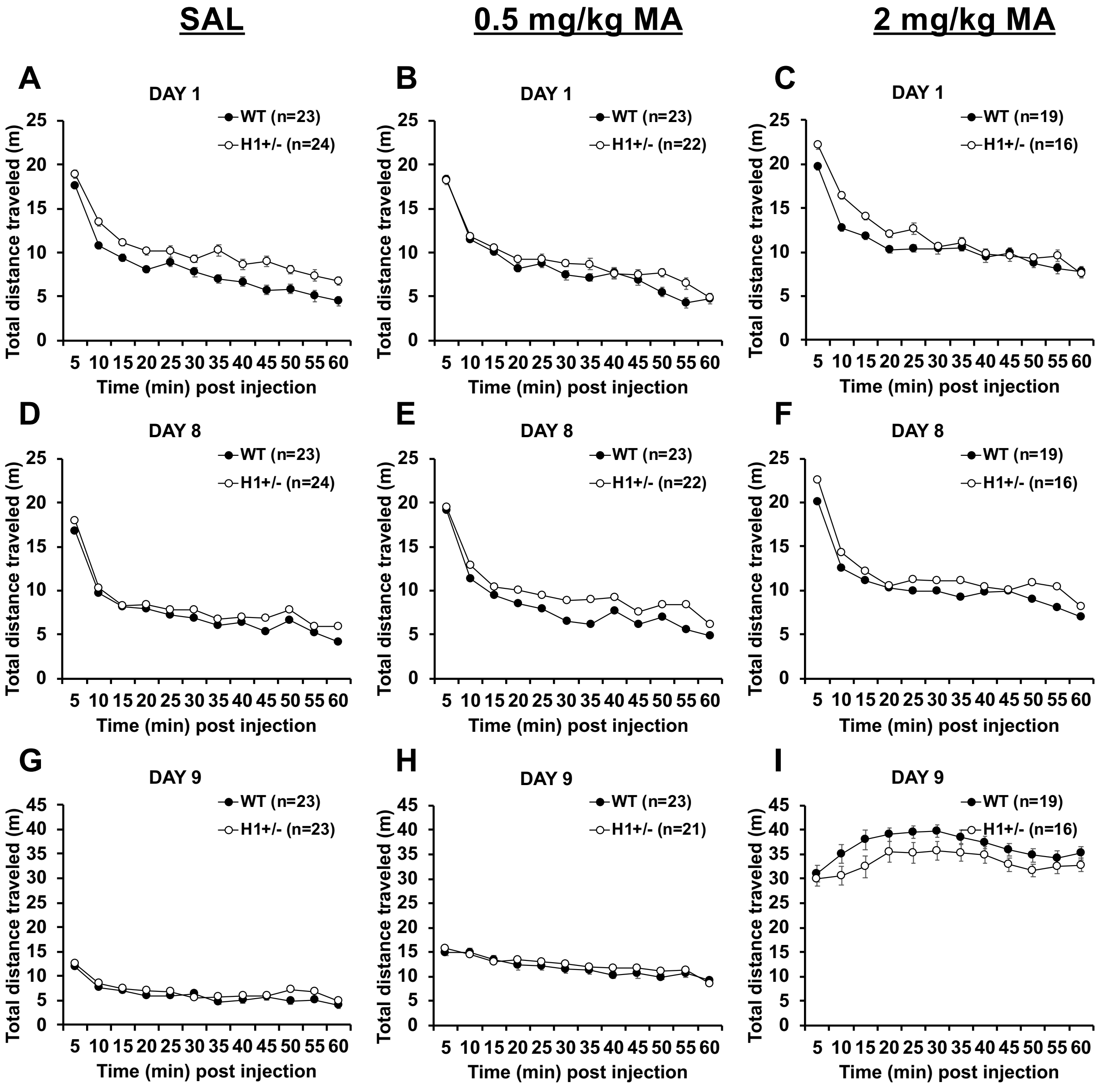
SAL
0.5 mg/kg MA
2 mg/kg MA
A
DAY 1
WT (n=23)
H1+/- (n=24)
B
DAY 1
WT (n=23)
H1+/- (n=22)
C
DAY 1
WT (n=19)
H1+/- (n=16)
D
DAY 8
WT (n=23)
H1+/- (n=24)
E
DAY 8
WT (n=23)
H1+/- (n=22)
F
DAY 8
WT (n=19)
H1+/- (n=16)
G
DAY 9
WT (n=23)
H1+/- (n=23)
H
DAY 9
WT (n=23)
H1+/- (n=21)
I
DAY 9
WT (n=19)
H1+/- (n=16)
Total distance traveled (m)
Time (min) post injection
5 10 15 20 25 30 35 40 45 50 55 60

### Figure S6

A

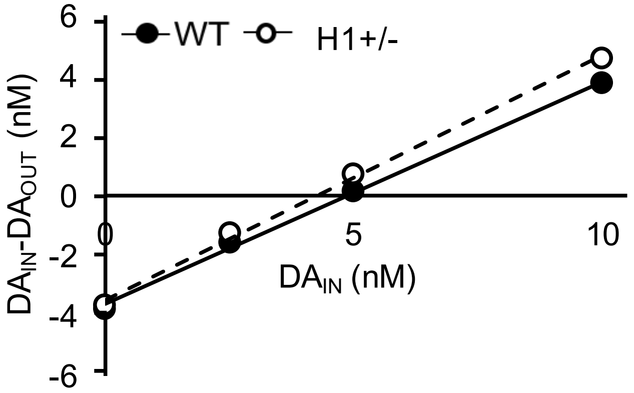


B

| WT | H1+/- |
|---|---|
| 0.70 ± 0.16 nM (n = 30) | 0.90 ± 0.17 nM (n = 30) |

C

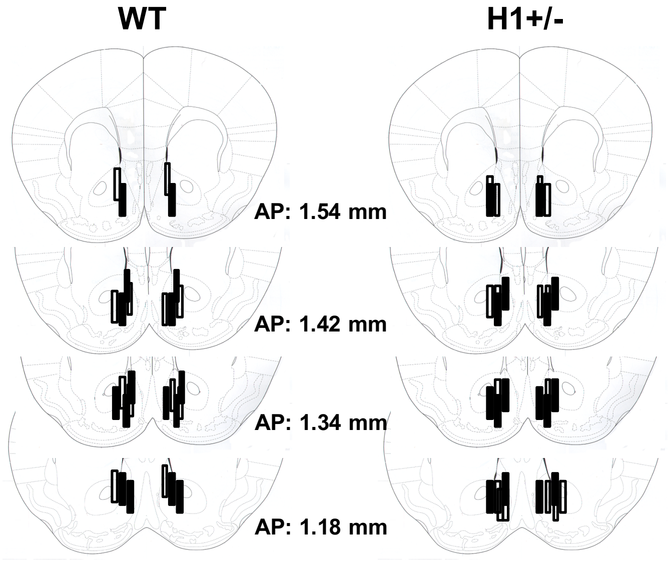

### Figure S7

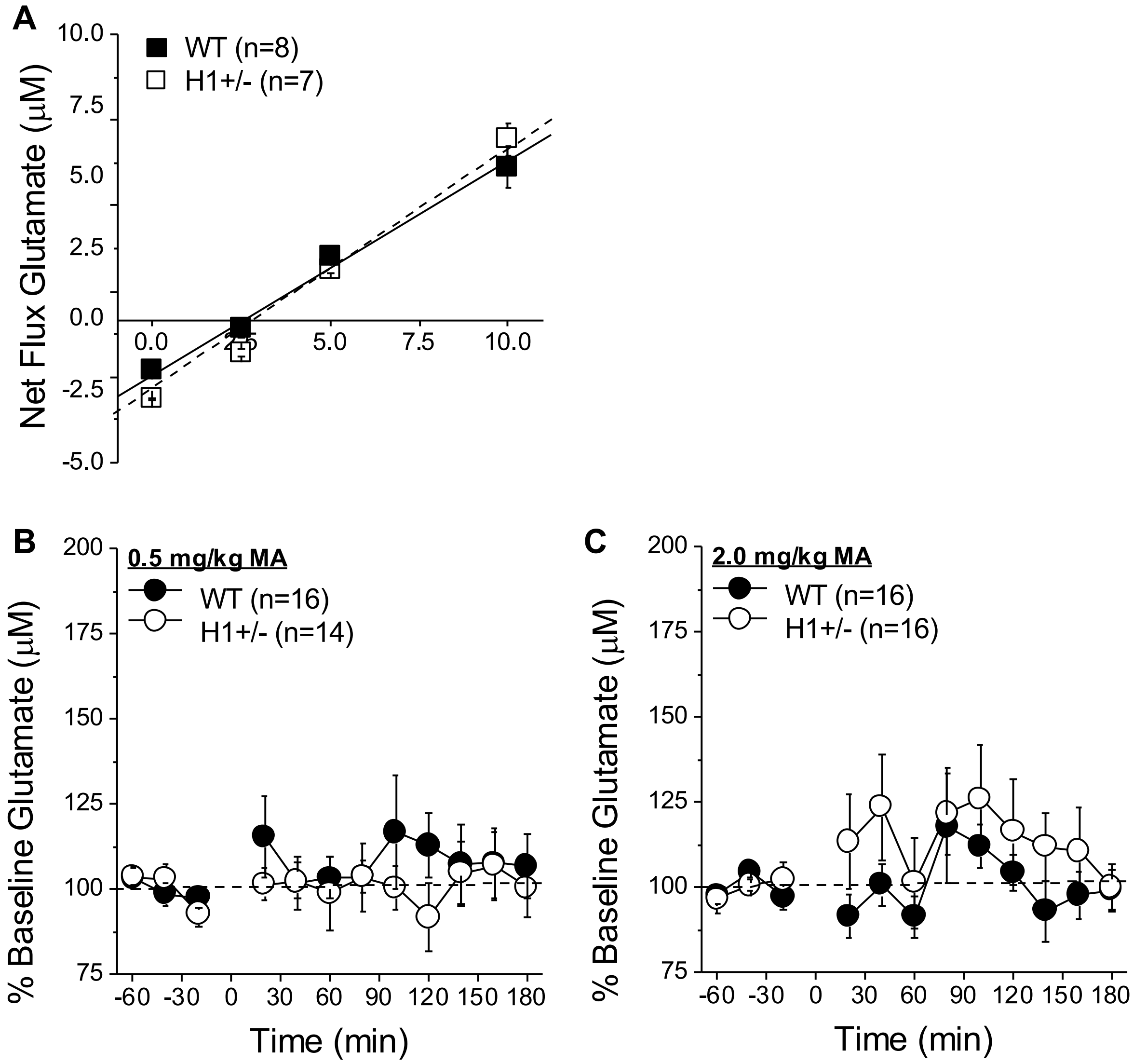
A
Net Flux Glutamate (μM)
WT (n=8)
H1+/- (n=7)
10.0
7.5
5.0
2.5
0.0
-2.5
-5.0
0.0
2.5
5.0
7.5
10.0
B
0.5 mg/kg MA
WT (n=16)
H1+/- (n=14)
% Baseline Glutamate (μM)
200
175
150
125
100
75
-60 -30 0 30 60 90 120 150 180
Time (min)
C
2.0 mg/kg MA
WT (n=16)
H1+/- (n=16)
% Baseline Glutamate (μM)
200
175
150
125
100
75
-60 -30 0 30 60 90 120 150 180
Time (min)

### Figure S8

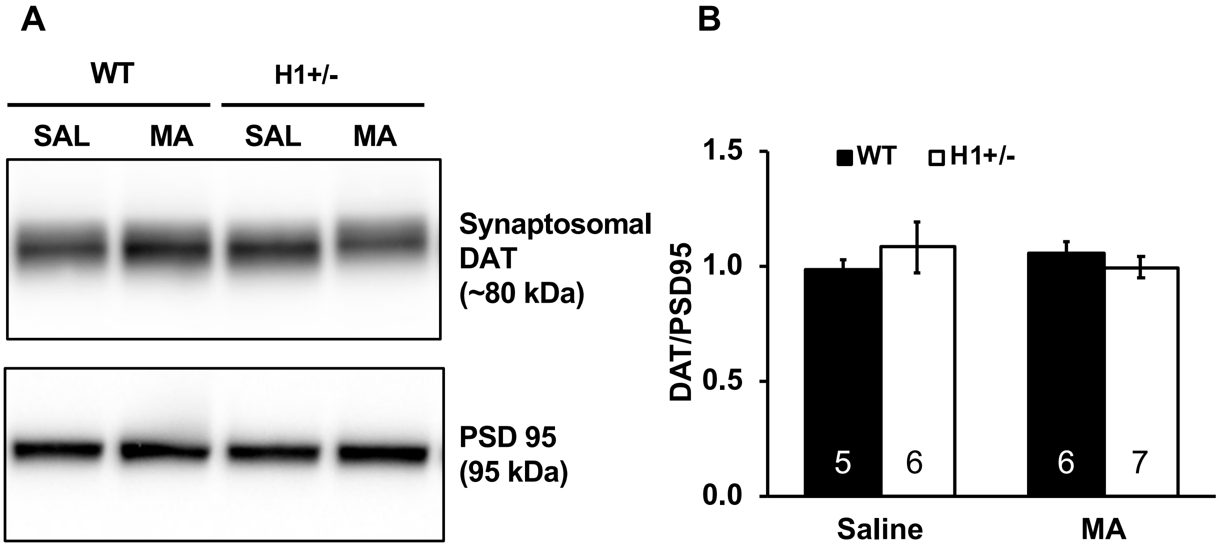
A
WT
H1+/-
SAL
MA
SAL
MA
Synaptosomal DAT (~80 kDa)
PSD 95 (95 kDa)
B
WT
H1+/-
DAT/PSD95
1.5
1.0
0.5
0.0
5
6
6
7
Saline
MA

### Figure S9

**A**

| Day | Assessments |
|---|---|
| 1 | Prepulse inhibition, Novel object |
| 2 | Marble burying, Light/dark, Porsolt Swim |
| 3 | Accelerating rotarod (3 trials) |
| 4 | Accelerating rotarod (3 trials) |
| 5 | Accelerating rotarod (4 trials) |

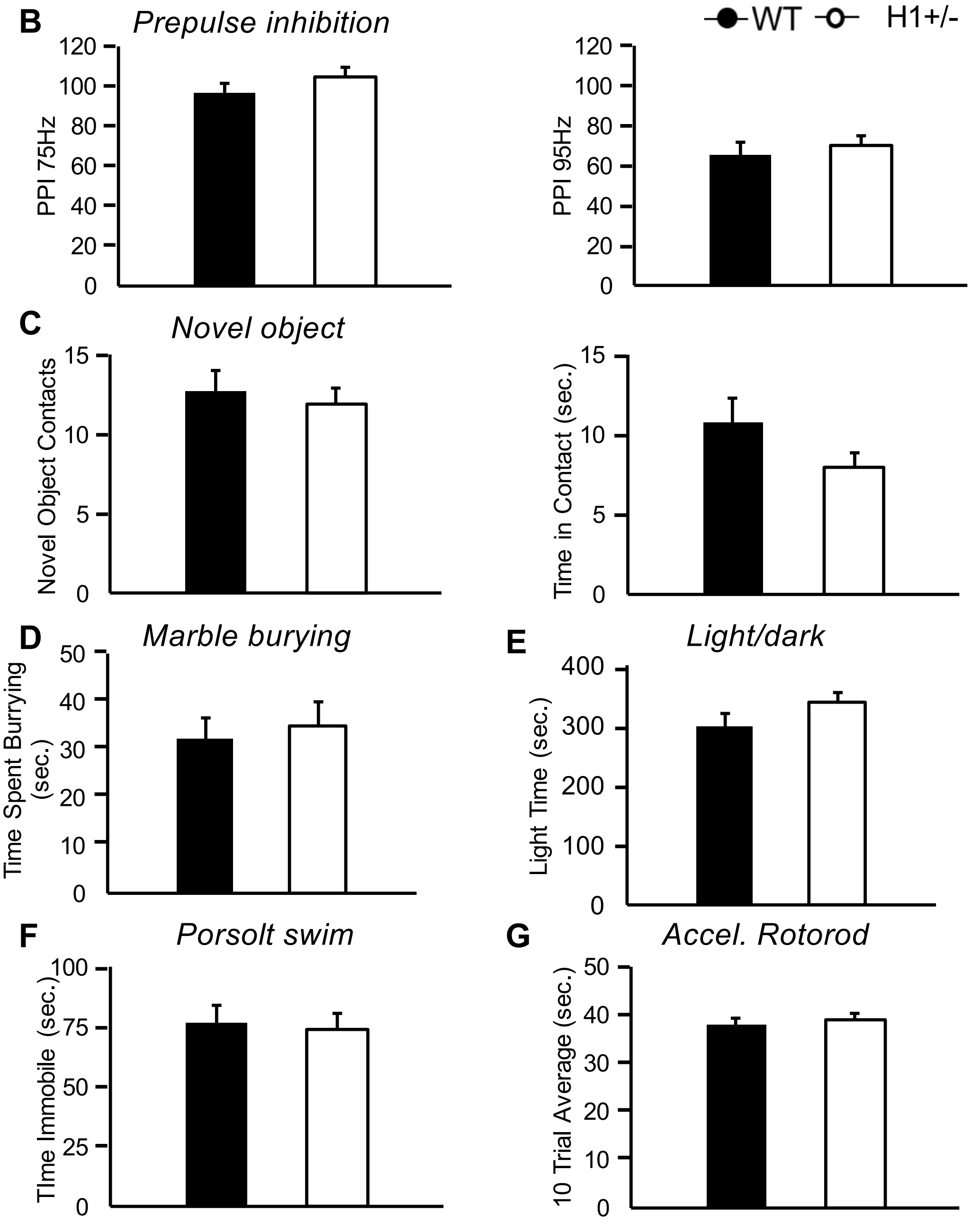

### Figure S10

A Bregma -3.28 mm to -3.64 mm

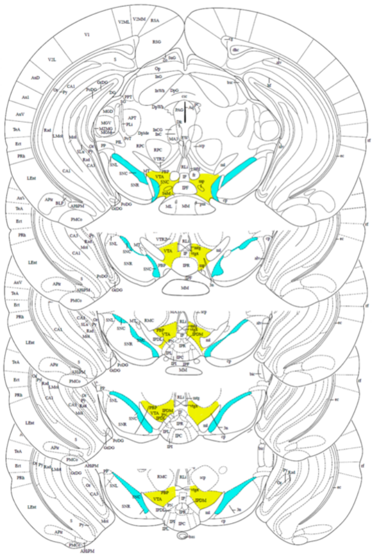

B Bregma 1.18 mm to 0.86 mm

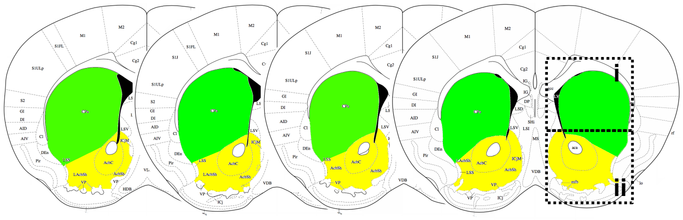

### Figure S11

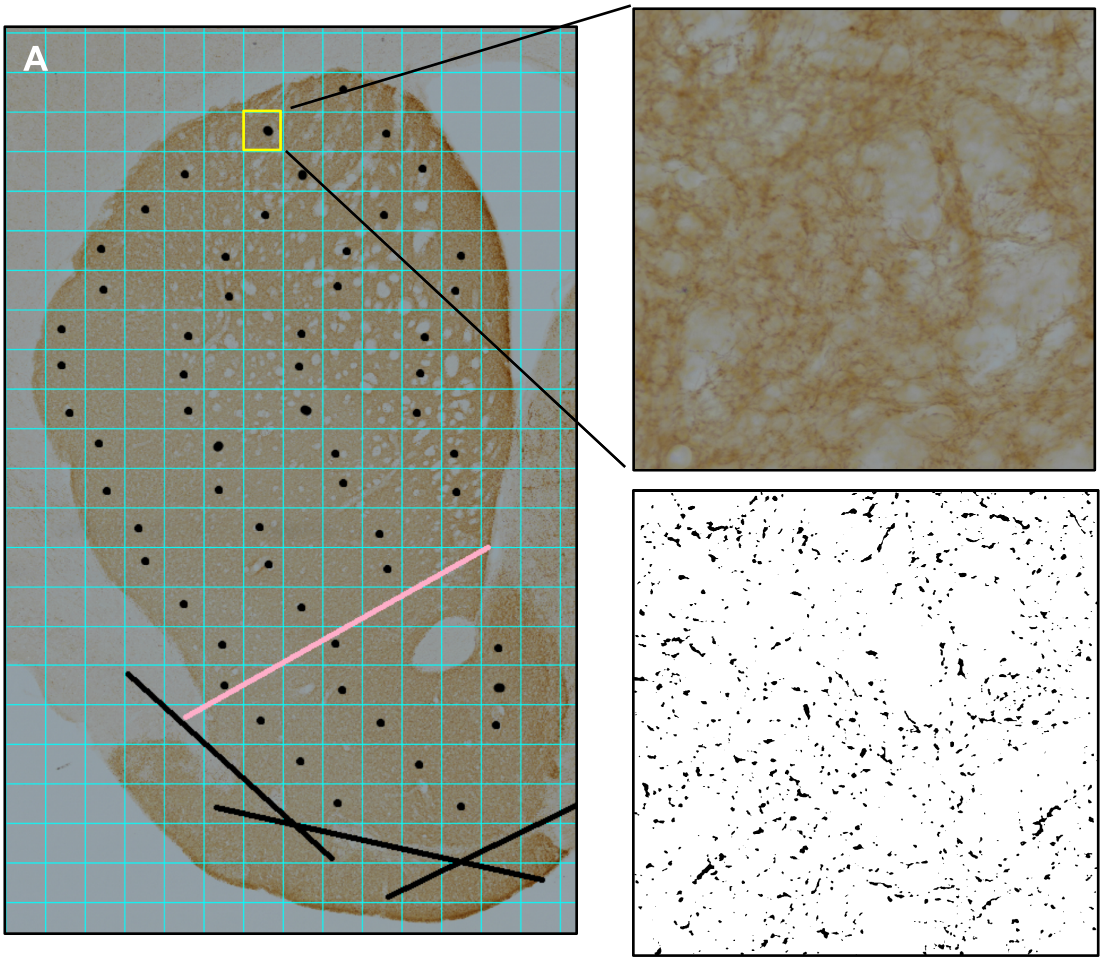
A

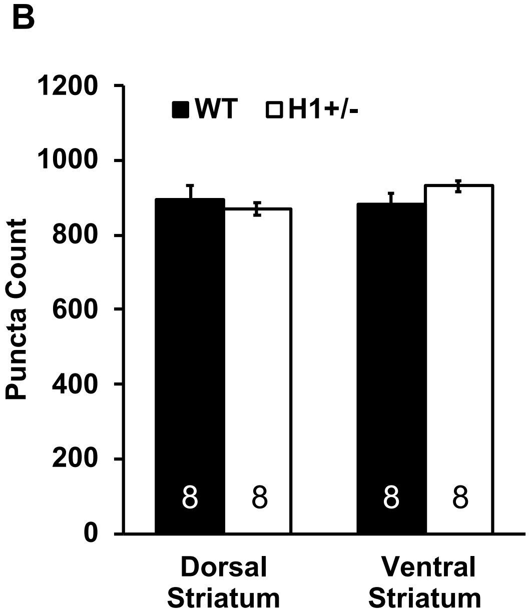
B
1200
1000
800
600
400
200
0
WT
H1+/-
Puncta Count
8
8
8
8
Dorsal Striatum
Ventral Striatum

### Figure S12

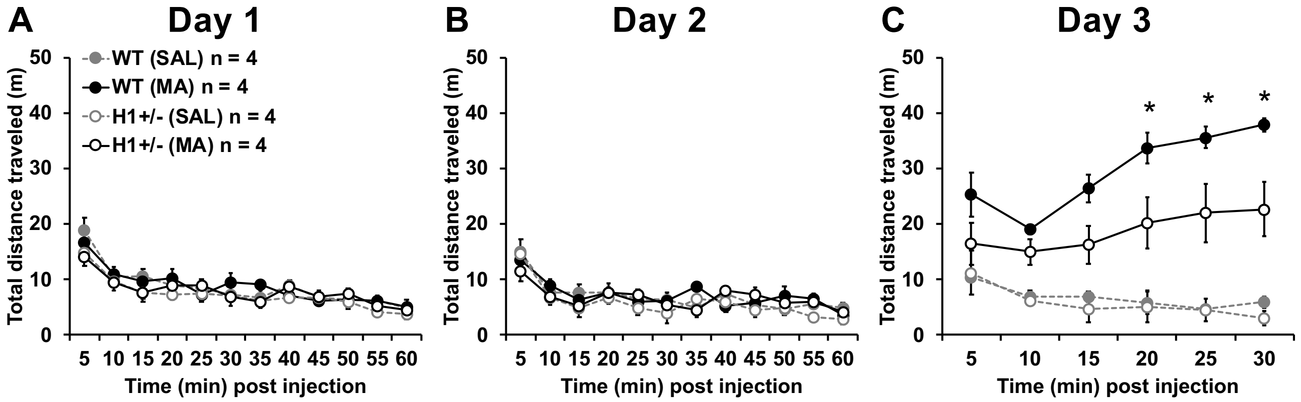
A
Day 1
B
Day 2
C
Day 3
WT (SAL) n = 4
WT (MA) n = 4
H1+/- (SAL) n = 4
H1+/- (MA) n = 4
Total distance traveled (m)
Time (min) post injection
*
*
*

#### Figure S13

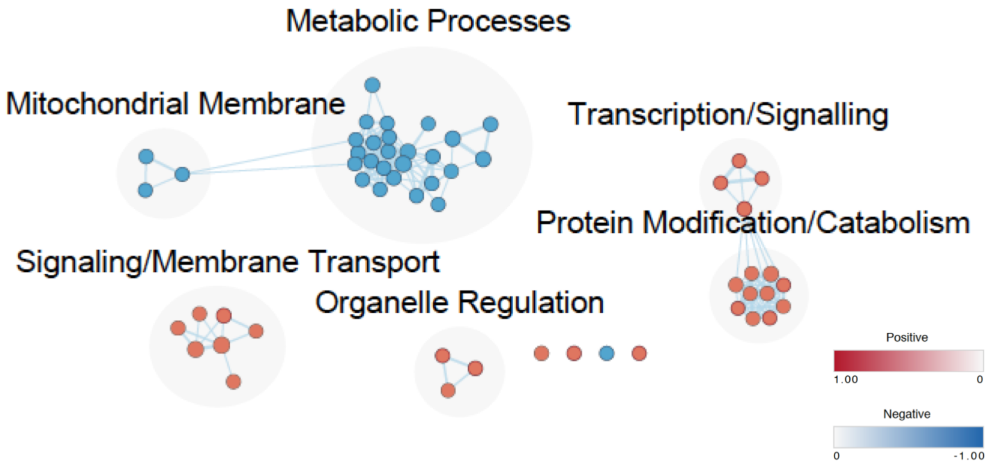
Metabolic Processes
Mitochondrial Membrane
Transcription/Signalling
Protein Modification/Catabolism
Signaling/Membrane Transport
Organelle Regulation
Positive
1.00
0
Negative
0
-1.00

### Figure S14

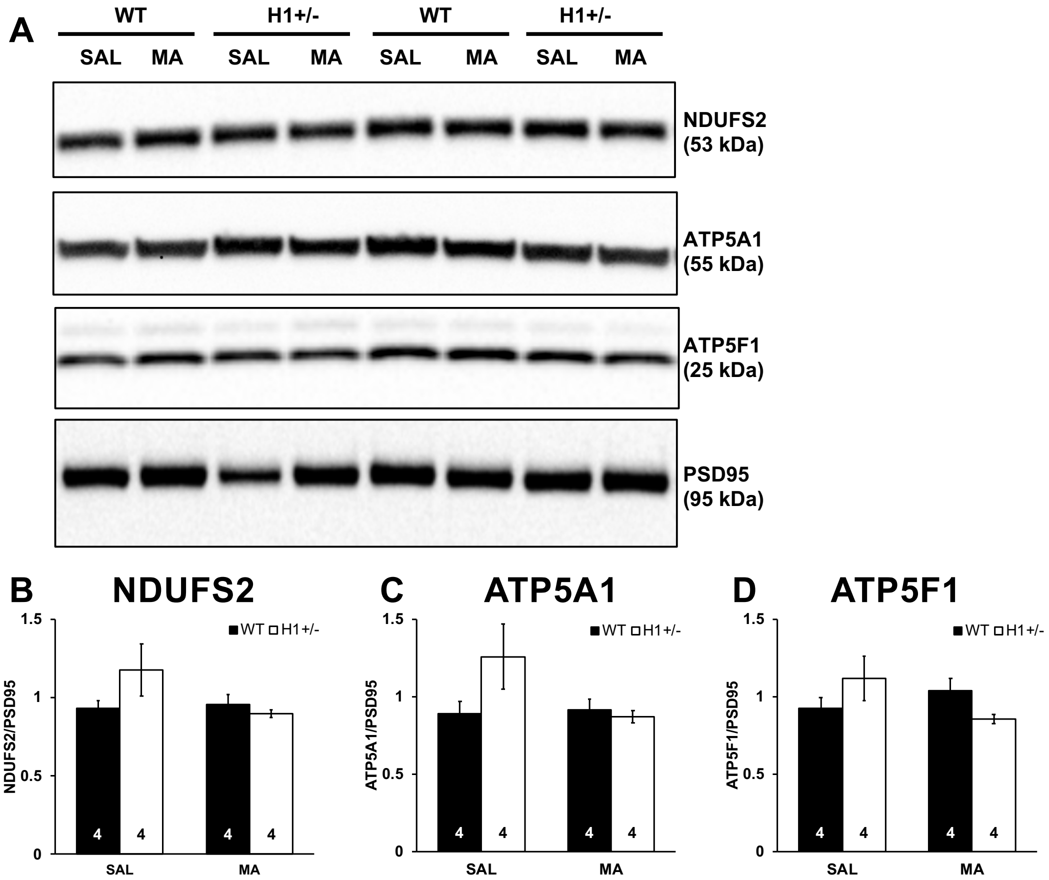
A
WT
H1+/-
WT
H1+/-
SAL
MA
SAL
MA
SAL
MA
SAL
MA
NDUFS2 (53 kDa)
ATP5A1 (55 kDa)
ATP5F1 (25 kDa)
PSD95 (95 kDa)
B
NDUFS2
NDUFS2/PSD95
WT
H1+/-
1.5
1
0.5
0
4
4
4
4
SAL
MA
C
ATP5A1
ATP5A1/PSD95
WT
H1+/-
1.5
1
0.5
0
4
4
4
4
SAL
MA
D
ATP5F1
ATP5F1/PSD95
WT
H1+/-
1.5
1
0.5
0
4
4
4
4
SAL
MA
